# Insight into the genetic aetiology of retinal detachment by combining small clinical and large population-based datasets

**DOI:** 10.1101/581165

**Authors:** Thibaud S. Boutin, David G. Charteris, Aman Chandra, Susan Campbell, Caroline Hayward, Archie Campbell, UK Biobank Eye & Vision Consortium, Priyanka Nandakumar, David Hinds, 23andMe Research Team, Danny Mitry, Veronique Vitart

**Author notes:** **Corresponding author:** Veronique Vitart.

## Abstract

Idiopathic retinal detachment is a serious common condition, but genetic studies to date have been hampered by the small size of the assembled cohorts. Genetic correlations between retinal detachment and high myopia or cataract operation were high, respectively 0.46 (SE=0.08) and 0.44 (SE=0.07), in the UK Biobank dataset and in line with known epidemiological associations. Meta-analysis of genome-wide association studies using UK Biobank retinal detachment cases (N=3977) and two cohorts, each comprising ∼1000 rhegmatogenous retinal detachment patients, uncovered 11 genome-wide significant association signals, near or within *ZC3H11B, BMP3, COL22A1, DLG5, PLCE1, EFEMP2, TYR, FAT3, TRIM29, COL2A1* and *LOXL1.* Replication in the 23andMe dataset, where retinal detachment is self-reported by participants, firmly establishes association at six loci *FAT3, COL22A1, TYR, BMP3, ZC3H11B* and *PLCE1.* The former two seem to particularly impact on retinal detachment, the latter three shed light on shared aetiologies with cataract, myopia and glaucoma.

**Author Summary:** Retinal detachments are common conditions that may lead to permanent severe sight reduction or blindness; they are a major cause of emergency eye surgery. The most common type of retinal detachment follows a break in the retina and is thought to be in part genetically determined but little is known about the contributing individual genetic risk variants. The condition prevalence increases with age and with common eye conditions such as myopia, cataract or glaucoma. We showed that the retinal detachment cases derived from self-report or hospitalisation records in the large UK Biobank dataset show very similar characteristics to samples of carefully clinically evaluated retinal detachment with break cases and therefore could be used to perform genetic analysis of the condition. Association studies require large sample of cases and by pooling Biobank and clinical cases, this study identifies 11 novel significant associations, six of which were further replicated in an independent population-based dataset (23andMe). Two of the replicated findings seem to specifically underline retinal detachment risk while three others highlight shared genetic risk with myopia, cataract and/or glaucoma, paving the way to better understanding of these conditions and of their overlap.

## Introduction

Rhegmatogenous retinal detachment (RRD) is a cause of emergency ophthalmic intervention. It is initiated by a full thickness break in the retina leading to invasion of vitreous humour between the neurosensory retina and the underlying nourishing retinal pigmented epithelium. This leads to blindness if untreated. The annual incidence of RRD in the U.K. has been estimated at 12 per 100,000 population [1]. Despite recent advances in surgical treatment, the reattachment success rate after one operation has levelled out at ∼80%[2,3], and retinal function remains suboptimal even after successful reattachment[4].

RRD has been well studied epidemiologically. Posterior vitreous detachment (PVD), a common condition following age-related liquefaction of the vitreous gel [5], is thought to be a frequent initiating event[6,7]. Trauma, cataract surgery, myopia and diabetic retinopathies can accelerate vitreous changes and have all been associated with RRD[6]. Prior cataract surgery has been reported in a fifth to a third of individuals in European RRD cohorts[8,9,10,11], with high myopia the strongest RRD increasing risk factor in those individuals (a six fold multi-adjusted hazard ratio), above the type of cataract surgery or history of trauma[12]. Myopia is also a prominent feature in ∼50% of cases who have never undergone cataract surgery[11,13] and RRD onset under the age of 50 years old is strongly associated with high myopia[8,12,14]. Retinal conditions such as lattice degeneration[15,16] may additionally predispose to the retina breaking (holes and tears). Finally, most epidemiological studies have also reported male gender to be at higher risk of RRD[10,11,12,17], even when trauma cases are excluded [12,17].

This common condition has been poorly characterised genetically due to the small sample sizes of collections assembled. Besides the Mendelian syndromic conditions associated with RRD[18,19], some genetic contribution to non-syndromic RRD has been suggested by familial aggregation studies, with roughly a doubling of lifetime risk in first degree relatives of cases compared to that of controls[20,21]. The proportion of RRD liability contributed to by common genetic variants was estimated at 27% in the very first RRD genome-wide association study (GWAS), which we performed using 867 Scottish cases[22]. Follow-up of the top associated variants in this GWAS identified a common coding variant in the CERS2 gene, rs267738, significantly associated with RRD. However, it is likely that many true positive signals were missed. Additionally, the reported genetic contribution and significant association of *CERS2* need to be replicated. Independent rare mutations in *COL2A1* (in which Stickler syndrome causal variants have been described) have been reported to co-segregate with autosomal dominant RRD in several families with no non-ocular systemic features[23,24], and a burden of rare variants in this gene as well as a common intronic variant, rs1635532, have recently been suggested to contribute to idiopathic RRD[25].

The UK Biobank is a major international research resource designed to help identify the genetic and non-genetic causes of complex diseases burdening middle and old age[26]. Data linkage to the UK National Health Service (NHS) hospital in–patients records have been achieved for a large portion of the 500,000 participants. This allows identification of retinal detachment (RD) cases with some additional details. RRD is the most common RD type, although other forms exist (tractional and the very rare exudative RD). In addition, to capture potential cases not linked to medical records, at recruitment, all UK Biobank participants completed a touchscreen questionnaire followed by verbal interviews including any history of retinal detachment. This ascertainment is *a priori* less precise than the hospital records linkage. However, RD, requiring surgical intervention, appears to be a significant ophthalmic event that has a memorable impact on patients [20] and for other conditions, self-report has been shown to be specific, while lacking in sensitivity[27,28].

We first evaluated RD based on self-report in the UK Biobank using the overlap with hospital admission records where possible, together with the wealth of other information gathered on participants, including data on high myopia and cataract operations. We performed a genome-wide association study (GWAS) in the UK Biobank using the union of self-reported and hospital record linked cases to maximise power in a sample of cases anticipated to be heterogeneous. To interpret the genetic associations uncovered, we carried out sensitivity analysis using phenotypic subsets in this discovery sample, as well as overlapping associations with high myopia and cataract, and tests of association in independent sets of clinically ascertained RRD cases from Scotland and England. Finally, we performed the largest GWAS analysis to date for retinal detachment by combining the UK-Biobank data with studies of clinically ascertained RRD and performed a look-up, in a population-based sample from the personal genetics company 23andMe, Inc. data, for the index variants from the genome-wide significant signals (study workflow summarised Fig S1).

## Results

### Evaluation of UK Biobank self-reported retinal detachment

1754 UK Biobank participants self-reported retinal detachment (RD-SR) at any one of the three assessment visits. Questionnaire responses were fairly consistent in the baseline visit, with 88% of RD-SR cases also indicating a condition which would prevent them from undergoing spirometry (compared to 7.7% in all respondents), 79.4% to having had an eye problem other than wearing glasses (compared to 14.7% in all respondents), and 79.8% having had a retinal operation/vitrectomy operation (compared to 0.6% in all UK Biobank participants).

RD-SR cases display significant distributional shifts, compared to controls (defined in S2 Text), for characteristics associated with RRD in epidemiological studies: an older age (median year of birth 1947 compared to 1951 in controls), a larger proportion of men (59.1% compared to 45.14%), higher prevalence of cataract operations, ocular trauma, diabetes (Fig 1). The proportion of glaucoma in the RD-SR was also very similar to that reported, 9.5%, amongst patients undergoing primary operation for retinal detachment[29]. The median age for first use of lens to correct for myopia was 12 year old in cases compared to 17 in controls. The self-report of age at first use of and reason for optical correction have been shown to accurately identify myopia, and age at first use to inversely correlate with severity of myopia [30]. The shift observed in RD-SR cases towards younger age of onset therefore suggests higher incidence of high myopia, a well-recognized risk factor for RRD. High myopia may also explain the increased incidence of glaucoma and macular degeneration in cases, as these are other potential pathological complications. The distribution shifts are not due to sex bias, as they are maintained in the same gender comparisons (Fig 1); although the expected sex difference in the proportion of eye trauma in SR cases was present with a higher frequency in men (P=0.012).

**Fig 1.**
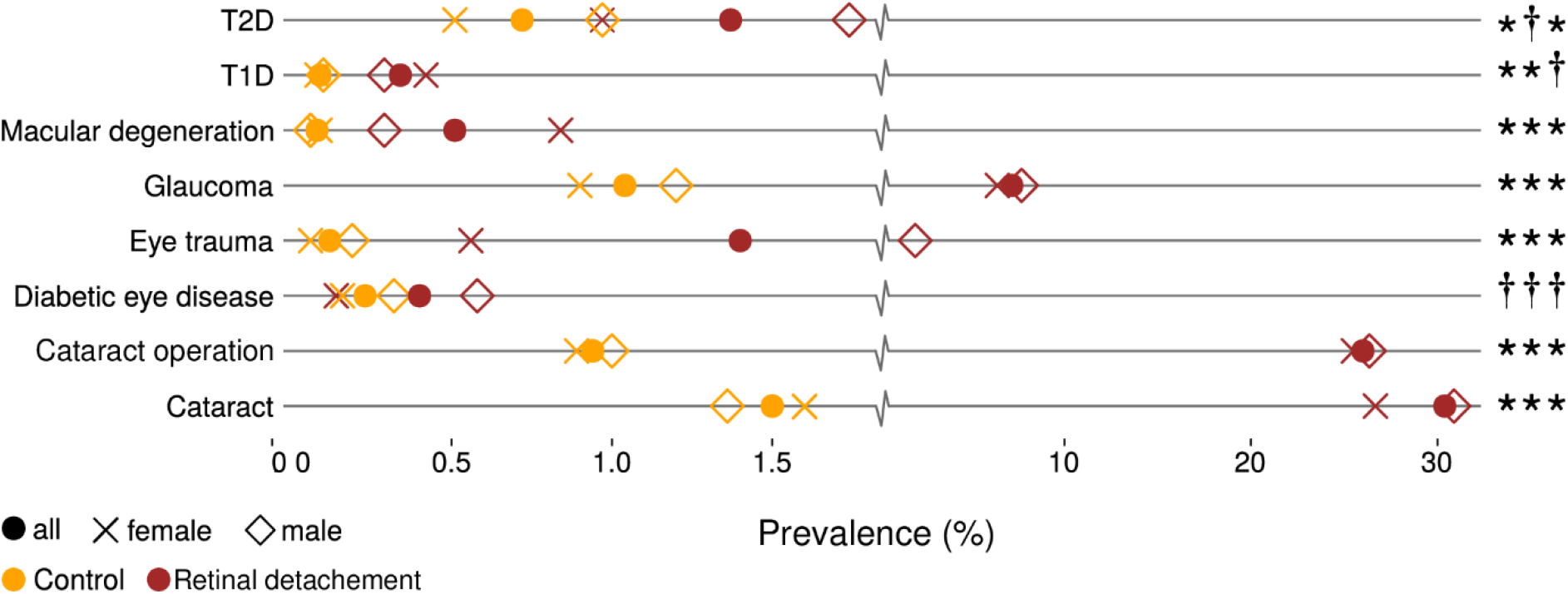
Comparison of self-reported features between self-reported retinal detachment cases and controls in UK Biobank. Across all ethnicities, a total of N=1754 UK Biobank participants self-reported a retinal detachment, leaving N=457,255 controls (see description in Supplemental note 2). * denotes a significant difference based on a two sided proportion z test at a 1% threshold; † denotes no significant difference. Symbols are reported respectively for the all, female and male comparisons. Symbols: T2D: Type2 diabetes, T1D: type 1 diabetes.

Linkage to retinal detachment in hospitalisation records was found for ∼half (N=890) of the RD-SR cases and shows that they potentially encompass a heterogeneous set of retinal conditions (Table 1). The highest proportion corresponded to RRD, ICD10 code H33.0 (retinal detachment with retinal break). The RD-SR cases that are linked to an H33.3 code (retinal break, no detachment) may reflect imprecise coding in hospital entries and/or that the breaks are a risk for future retinal detachment. Fig 2 displays retinal detachment ICD9 and ICD10 entries in the overall UK Biobank participants and illustrates the extent of common overlaps between the different sub-codes.

**Fig 2.**
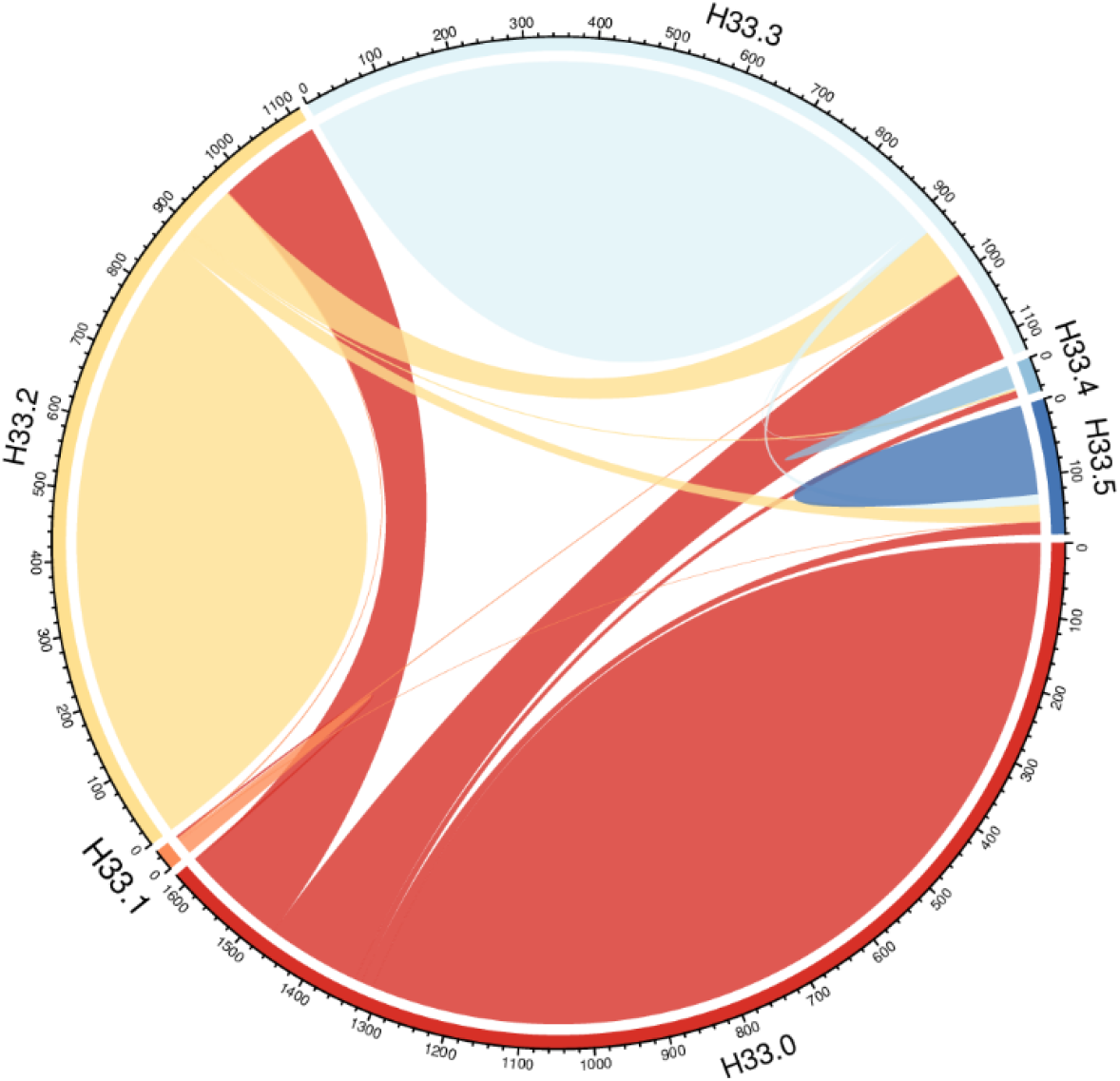
Distribution of “retinal detachments and breaks” sub-types amongst UK-Biobank participants linked to an ICD10 code H33. H33 encompasses subclasses H33.0 (retinal detachment with retinal break), H33.1 (retinoschisis and retinal cysts), H33.2 (serous retinal detachment), H33.3 (retinal breaks without detachment), H33.4 (traction detachment of the retina) and H33.5 (other retinal detachments). The numbers represent the UK Biobank participants (all ethnicities) associated with each code. The plot was drawn with the R package *circlize* v0.4.3[75].

**Table 1.**
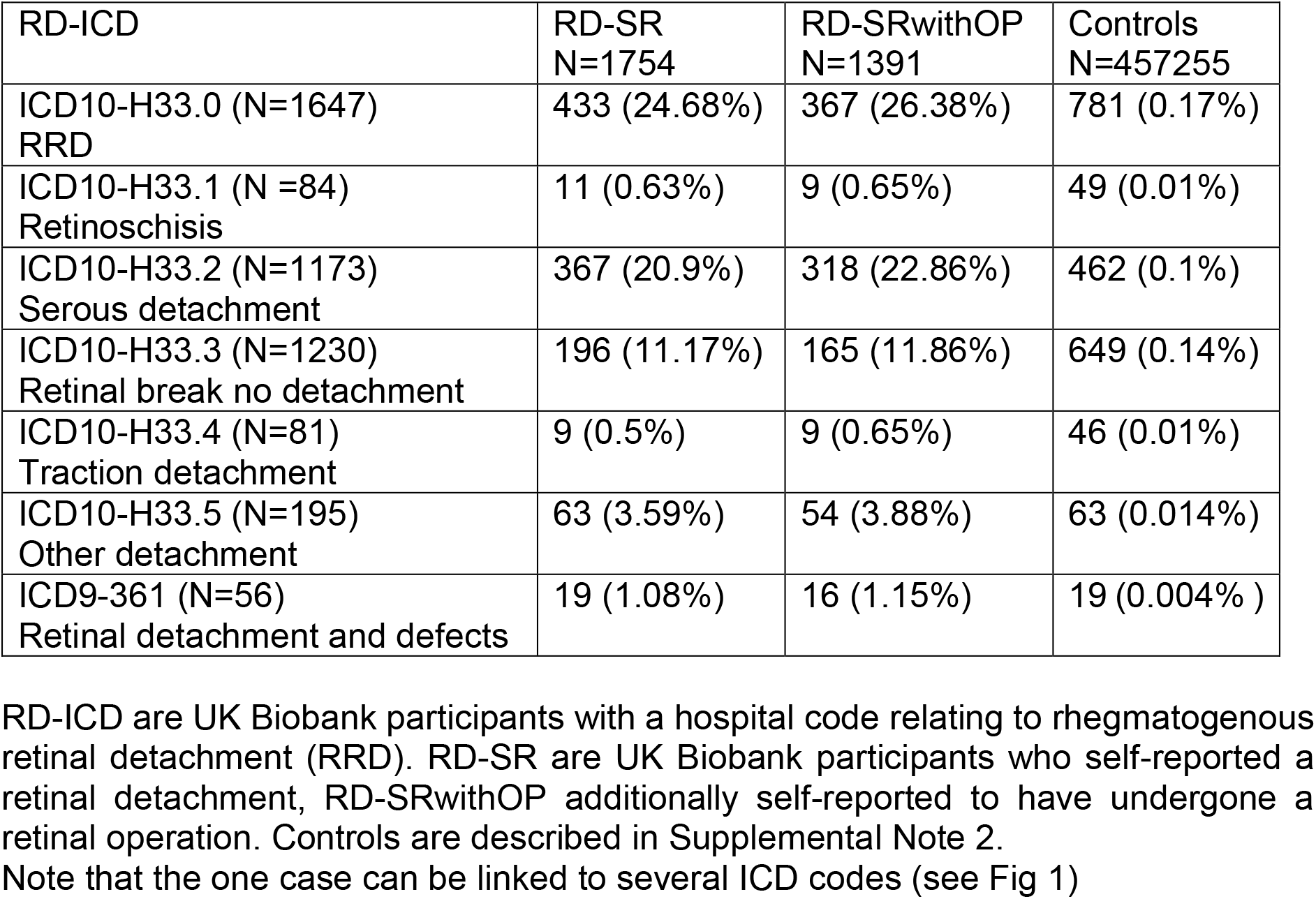
Overlap between self-reported and hospitalisation records in UK Biobank

There seems little gain in restricting RD-SR cases to the individuals who self-report a retinal operation/vitrectomy for greater accuracy: 363 cases would be excluded, but these do not change the overall characteristics of the RD-SR cohort considered (Table 1). Given the enrichment in RD cases that they are likely to represent, we combined all SR cases with individuals linked to the ICD10 super code H33 and ICD9 code 361 to increase power in the initial association study (RD-SR-ICD).

### Genetic variants enriched in the UK Biobank SR and ICD combined retinal detachment cases

GWAS identified two genome-wide significant signals with common lead variants rs28531929 (MAF=31.9%, P=4.6×10^−10^) and Affx-6370160 (rs10765565; MAF=39.8%, P=1.7×10^−13^) in the genotyped dataset, on chromosomes 8 and 11 respectively. Using the imputed dataset, these signals were refined respectively, to rs11992725 (MAF=31.9%, P=3.1×10^−10^), an intronic variant in *COL22A1*, and rs7118363 (MAF=39.2% P=1.2×10^−16^), an intronic variant in *FAT3*. Analysis using the imputed dataset revealed a third genome-wide significant locus with lead variant rs633918 (MAF=38%, P=1.6×10^−8^), an intronic variant in *GRM5* (Fig S2 and regional plots Fig 3A).

**Fig 3.**
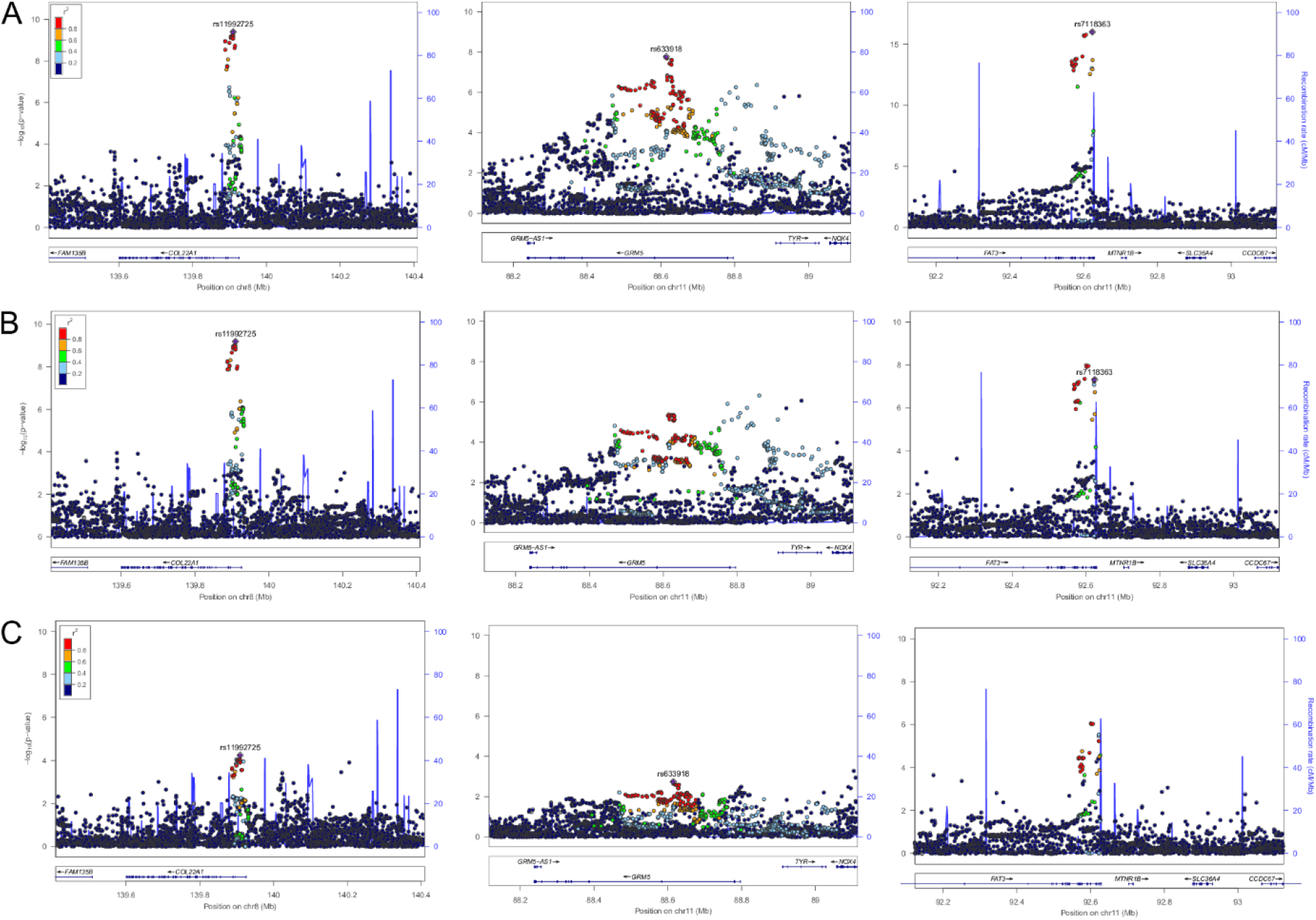
Regional association plots at the three genome-wide significant loci in the primary UK Biobank retinal detachment analysis across different case definitions. The y axis represents the negative logarithm (base 10) of association P-value and the x axis the position on the chromosome. The index variants from the RD-SR-ICD discovery study (A – Ncases=3977) are marked by a purple diamond. The colours of the other variants reflect LD strength (r^2^) with the index variant at each locus. (B): results at the same loci when cases linked to ICD10 codes H33.1 and H33.3 are omitted (Ncases=2893). (C): results when cases are restricted to participants with H33.0 code (Ncases=1380).

No other strong disease/trait associations have previously been reported with the *COL22A1* and *FAT3* loci index variants (or with 1000G proxy variants in high LD, r^2^>0.6). In Phenoscanner [31], the top associated phenotypes, as of May 2018, were waist circumference for the *FAT3* locus (proxy rs7101614, P=5.8×10^−5^, N=145346, GIANT consortium) and hip circumference for the *COL22A1* locus (proxy rs2318346, P=3.36×10^−3^, N=85539, GIANT consortium). PheWAS from the Global Biobank Engine (GBE; URL: http://gbe.standford.edu/) or Gene Atlas (http://geneatlas.roslin.ed.ac.uk/) servers also indicate specificity for both signals amongst UK Biobank traits. In contrast, the index variant in *GRM5* is in high LD with variants that have been strongly associated with eye colour (e.g proxy rs7120151 P=2.45×10^−10^)[32]. In GBE, the top associated phenotypes are sunburn sensitivity and skin cancer, with many phenotypes displaying high to moderate level of association including intra ocular pressure and cataract.

A full list of loci with lead variants displaying an association P < 5×10^−6^ is shown in S1 Table. Two of the 24 suggestive loci (5×10^−6^<P≤5×10^−8^) are amongst the most strongly supported loci associated with common myopia [33]: LAMA2 (identical lead variant) and BMP3 (reported myopia lead variant rs5022942 [33] in some linkage disequilibrium, r2= 0.19, D’=0.999, with the RD suggestive locus index variant).

The quantile-quantile plot for the imputed dataset GWAS results (S2 Fig-λ_GC_=1.097) and the LDscore regression intercept of 1.0083 (0.0067) suggest an excess of genetic associations attributable to polygenic effects, compared to the null expectation of no genetic effect. Using a more balanced, 1:3, case:control design, that has been advocated [34] to avoid possible distortion of results for rare variants (which are not analysed here), yields the same top two genome-wide significant signals but with less significant P-values (S3 Fig), indicating some loss of power when using a reduced set of controls.

The *COL2A1* common variant rs1635532 previously proposed to be associated with RRD [25] displays no association (P=0.95) in the UK Biobank, but other variants in *COL2A1* which is an excellent candidate gene [24] displayed a suggestive level of association, with P=1.5×10^−6^ for the genotyped variant rs1793959. The *CERS2* variant rs267738 previously associated with RRD[22] shows nominally significant association, P-val=4.2 10^−3^, with the major T allele increasing risk as previously reported.

Jointly, the common variants tested contribute a heritability on the liability scale, 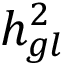, of 0.23 (P= 2×10^−72^). 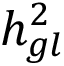 calculated using LD score regression was similarly low but significant, 0.1378 (SE=0.0194).

### Retinal detachment sub-analyses and genetic overlaps with high myopia and cataract in the UK Biobank dataset

To better understand what these genetic signals relate to, we carried out sensitivity analyses to tease out phenotypic heterogeneity. Performing the same GWAS analyses after removing cases linked to an ICD code suggesting a break in the retina but no detachment (ICD10 codes H33.3 and H33.1, leaving N=2893 cases of white-British ancestry) leads to a decrease in the *GRM5* signal in line with the reduction of cases (rs633918 P=4.6 x10^−6^), and a more marked decrease of the *FAT3* signal (top genotyped variant rs7118248 P=7.7×10^−8^, top imputed variant rs561742998 P=9.5×10^−9^) than expected. Of note, the *COL22A1* signal shows little change (top genotyped variant rs28531929 P=1×10^−9^, top imputed variant rs11992725 P=6.5×10^−10^) (Fig 3B), indicating little contribution to this signal from the cases linked to a H33.3 or 33.1 code. Two new signals reached genome-wide significant level in this sub-analysis, both driven by low frequency variants: *PDE4D* on chr5 with intronic variant rs570334020 (MAF 1%; P=3×10^−8^) and *CPAMD8* on chr18 with intronic variant rs192943660 (MAF=1.3%; P=4.3×10^−8^).

Performing GWAS in the much reduced set comprising only the cases linked to an H33.0 ICD code (Ncases=1380) – which should correspond most closely to RRD cases – does not yield any genome-wide significant associations. However, *FAT3* remains amongst the strongest signals (with P=4×10^−6^ for genotyped rs7118248, and P=8.6×10^−7^ for the top imputed variant rs10765567) (Fig 3C). Three loci in addition to *FAT3* have an index SNP with P < 10^−6^ in the imputed dataset (S2 Table), two of which have low minor allele frequency and are therefore prone to chance association given the low number of cases.

In addition to sensitivity analyses, we looked for suggestion in UK Biobank of shared signals with the two epidemiologically established conditions conferring increased retinal detachment risk: cataract surgery and high myopia, for neither of which a well powered GWAS has yet been published. GWAS analyses for high myopia (N=2737 cases) and history of cataract operation (N=21679 cases) showed contribution of common variants to risk, especially strong for high myopia, with 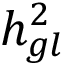 of respectively 0.723 and 0.125 (0.7217 (SE=0.0642) and 0.0719 (SE=0.0072) using the LDscore regression method). Manhattan plots for these analyses are shown in S4A-B Fig, and the lists of significantly associated loci in S3-S4 Table. In addition to the top associated locus, intergenic GOLGA8B-*GJD2* (index rs524952, P=1.2×10^−24^), nine out of the other sixteen high myopia loci show nominal association with RD-SR-ICD (P_RD_ < 0.05). All ten associations show agreement with the epidemiological association (allele increasing high myopia risk increases risk of retinal detachment). Two out of the 20 cataract operation associated loci show nominally significant association with RD, all with the cataract risk allele increasing risk of retinal detachment. These two loci, interestingly, encompass the eye colour genes *OCA2* and *NPLOC4*. The regional high myopia and cataract association plots for the three genome-wide significant retinal detachment loci *COL22A1*, *FAT3* and *GRM5* are shown in S5 Fig and suggest that those are not driven by a high myopia or cataract association. The look up for association values with high myopia and cataract for all the suggestive RD loci (P < 5×10^−6^ in the RD-SR-ICD analysis) is displayed in S5 Table. The *COL22A1* index variant shows nominal association with high myopia and cataract. Four additional loci, show nominally significant risk association across all three phenotypes in line with expectation from their epidemiological associations. The *FAT3* lead variant does show nominally significant association with high myopia (P=0.0069), but the effect is not direction consistent with the epidemiological association (the variant increasing retinal detachment risk protecting against high myopia).

In agreement with the look-ups, the overall genetic correlations calculated using LD score regression slopes [35] were high, 0.46 (SE=0.08) between RD and high myopia, 0.44 (SE=0.07) between RD and cataract, and 0.25 (SE=0.066) between high myopia and cataract, supporting substantial shared genetic aetiology.

### Comparison of RD genetic associations in UK Biobank and in datasets of clinically ascertained cases

Our findings in the UK Biobank were evaluated in two independent datasets, each comprising ∼1000 clinically ascertained RRD cases, collected in Scottish vitreoretinal surgeries or at the Moorfields London Hospital respectively. We performed GWAS scans using these clinically ascertained cohorts and population matched controls, using genotypes imputed to the HRC reference panel. Seven of the eight RD-SR-ICD lead variants with significant or suggestive (P ≤ 10^−6^) associations passed stringent quality controls (data clean up S4 and S5 Text) and their associations in RRD datasets shown Table 2. Using a significant replication P-value threshold of 7×10^−3^ to account for the seven look-ups (α=0.05/7), the *FAT3* signal replicates in both independent clinical RRD sets, with consistent effect direction and magnitude across studies. Index variants for the two other genome-wide significant loci from the UK Biobank analysis, and for the myopia locus *LAMA2*, all displayed nominal association in one of the two RRD cohorts but no evidence at all in the other. Variants for the two RD-SR-ICD suggestive loci, *BMP3* and *DLG5*, showed homogeneous effects across all studies but the P values do not reach the significance threshold for replication in both RRD studies, indicating limited power. The converse look up for the top 12 lead variants associated with RRD (P ≤ 5×10^−6^) after meta-analysis of the two clinic-based studies (S6 Table) indicates that in addition to *BMP3*, two other loci show nominal association in the UK Biobank analysis, 5’*ZC3H11B* and *UPP2*. Index variant in the former is in high LD with index variant of a newly reported refractive error locus[36], *LYPLAL1*. Summary statistics from the latest, large-scale, refractive error GWAS[36] was used to derive a simple aggregate myopia genetic risk score. The score distributions showed similar shift between cases and controls in both the UK Biobank study and the combined dataset of clinically ascertained RRD, with proportion of cases versus controls increasing with higher decile of the myopia risk score (S6 Fig). The odds for retinal detachment with increasing decile of the myopia scores, relative to that of the population most common decile, were significant and of similar magnitudes using RD defined in UK Biobank and RRD cases who had been clinically ascertained (Fig 4). Given these evidences, a GWAS meta-analysis for all studies (GWAMA) was performed. Of note, the *CERS2* variant rs267738 we previously reported as associated with RRD [22] did not show even nominal association in the London-based dataset (which only partially overlaps with the London-based dataset used previously [22]), despite good quality call for a genotyped variant in high LD with it, rs267734, on the Illumina GSA chip used.

**Fig 4.**
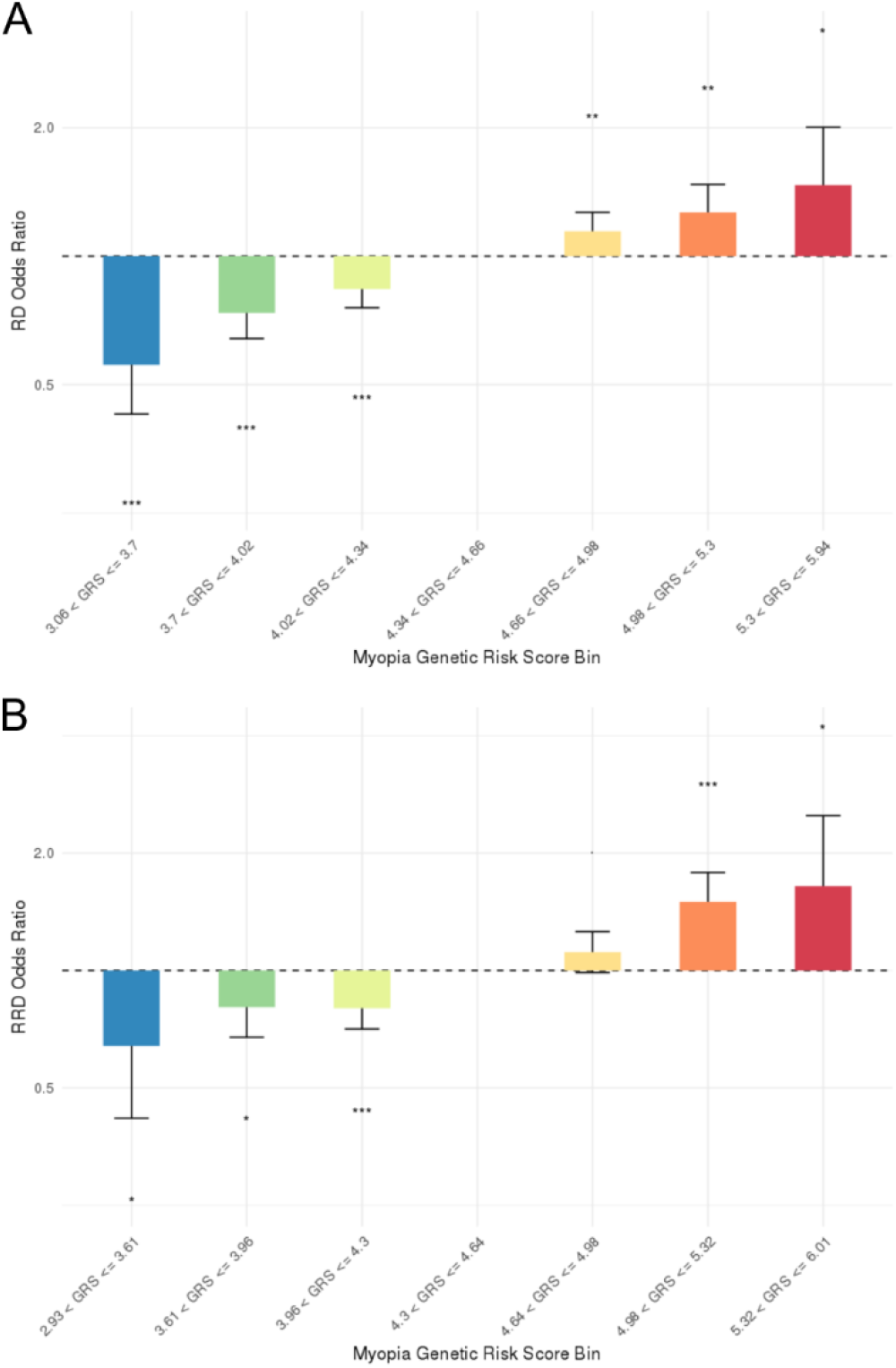
Myopia genetic risk score effect on retinal detachment risk using UK Biobank RD-SR-ICD cases or clinically ascertained RRD cases. The analysis of the myopia GRS on retinal detachment (RD) risk in UK Biobank and on Rhegmatogenous RD (RRD) in the aggregated clinically ascertained sets are displayed in A and B respectively. Odd-ratio for detachment is plotted along the y axis for separate decile of the GRS using the most common value in the UK population (S6 Fig) as the reference, the last two deciles of myopia GRS at each extreme were pooled to insure sufficient number of observations for testing. ‘***’ denotes a P-value reaching a significance threshold of 0.001, ‘**’ of 0.01 and ‘*’ of 0.05.

**Table 2.**
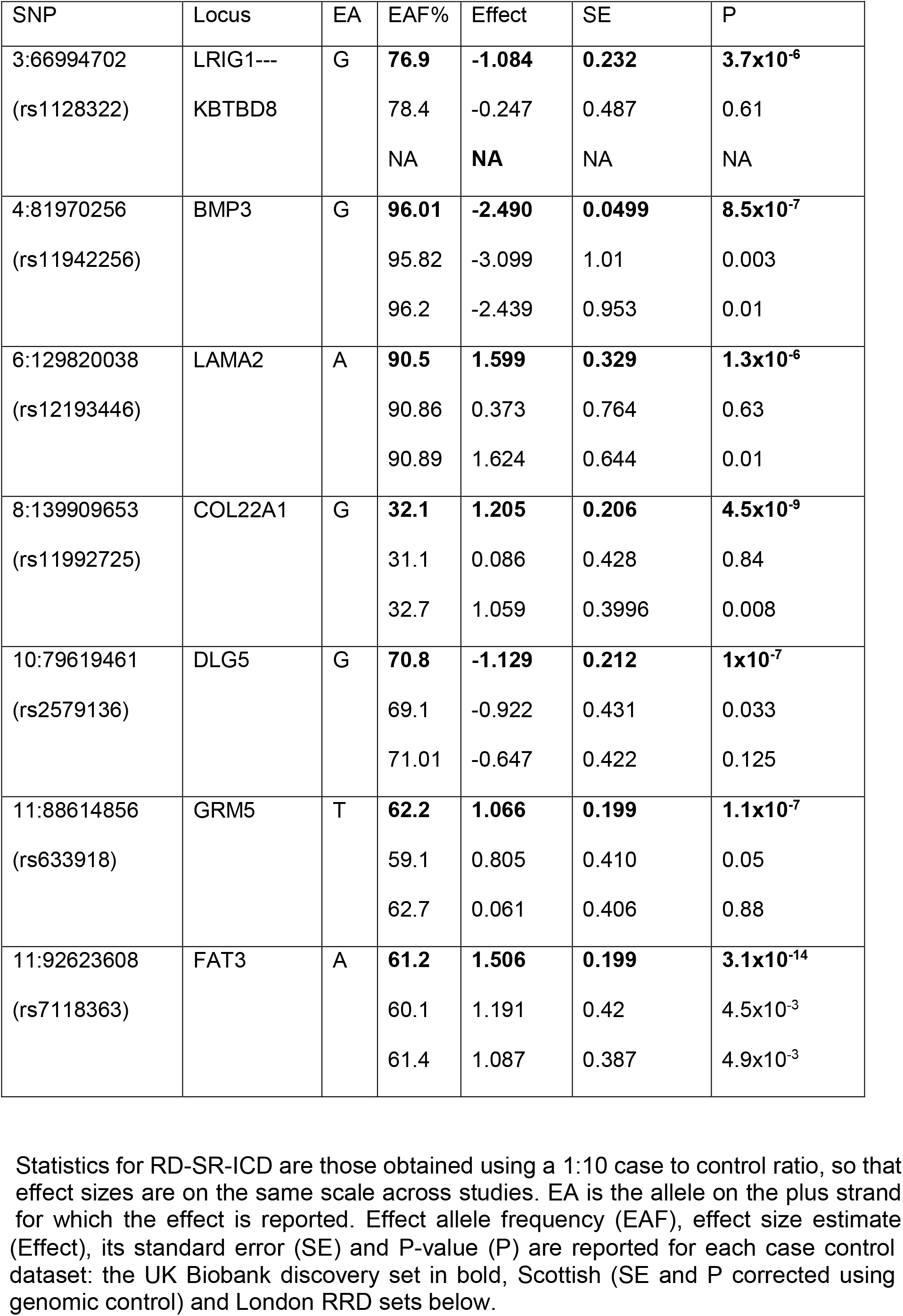
Look-up in the RRD GWAS of index variants associated at a P value cut-off of 10^−6^ in the UK Biobank retinal detachment study (RD-SR-ICD)

### GWAMA of UK Biobank and clinical retinal detachment datasets and replication in 23andMe dataset

Following GWAMA and removal of variants with high heterogeneity across studies, 241 variants distributed across 11 independent loci showed significant (P < 5×10^−8^) association with retinal detachment (Fig 5 and S7 Table). These loci include *FAT3*, *COL22A1*, *BMP3*, 5’*ZC3H11B* and *DLG5* but also, and with high homogeneity of effect across studies (Forest plot S7 Fig), *TYR* in place of the physically close *GRM5*. The lead variant in *TYR* is the common missense variant (TYR S192Y), with the A derived allele strongly associated with light skin colour [37] and absence of freckles [38] increasing the risk of retinal detachment in all three studies. In addition, the 5’ *COL2A1* locus, upstream of the already discussed excellent candidate RRD gene *COL2A1*, the 5’*EFEMP2* locus, *PLCE1*, an intergenic region between the *TRIM29* and *OAF* genes and *LOXL1* are also genome-wide significant. The *LOXL1* lead SNP is in high LD with the pseudo exfoliative glaucoma (PXG) strong effect variant, rs2165241, (r2=0.9, D’=1 in 1000G-GRB reference haplotypes), with the T allele increasing PXG risk in Caucasians [39] associated with reduced RD risk (P = 6.7×10^−7^). The Functional Mapping and Annotation of the genetic association (FUMA) tool [40] identifies 564 either independent significant SNPs or tagged variants. Those with distinctive functional annotations are listed in S8 Table and previously reported associations or top associations in UK Biobank PheWAS repositories for the lead SNPs are listed in S9 Table. Four loci harbour variants with a predicted deleterious effect amongst the top 1% pre-computed scores for over 8 billion variants (scaled CADD score > 20) [41]: the two missense lead variants in *BMP3* and *TYR* as well as a variant in *TRIM29* (rs12792820) and *ZC3H11B* (rs6682581). Maximal CADD scores in *FAT3* and *LOXLI* were also high (> 14). Evidence of potential transcriptional regulatory attributes was also evident in each of the 11 loci (S8 Table). In addition to the already acknowledged associations to myopia, refractive error or PXG, the annotations highlighted ocular axial length (and height) for the lead variant in 5’ *ZC3H11B* and vertical to cup optic disk ratio (a glaucoma endophenotype) for the *PLCE1* lead variant. The gene-based association analysis (S10 Table) pointed to four additional associated loci, *DIRC3*, *CMSS1*, *C16orf45* and *BMP2*. Transcriptional expression in human eye tissues (S11 Table) shows that many implicated genes are highly expressed in the retina (neuronal retina or retinal pigmented epithelium). Mouse mutations associated with abnormal retinal morphologies have been described in four genes, located at independent loci: *TYR*, *FAT3*, *C4orf22* and *COL2A1* (S11 Table). Rare functional variants in the latter, *COL2A1*, have been associated in humans with RRD with and without systemic Stickler syndrome phenotypes[23,24]. In the gene-rich fifth locus, 5’*EFEMP2*, human loss of function variants in *EFEMP2* have been associated with cutis laxa, a condition where the skin lacks elasticity, with retinal pigmentation a commonly associated feature (https://rarediseases.info.nih.gov/).

**Fig 5.**
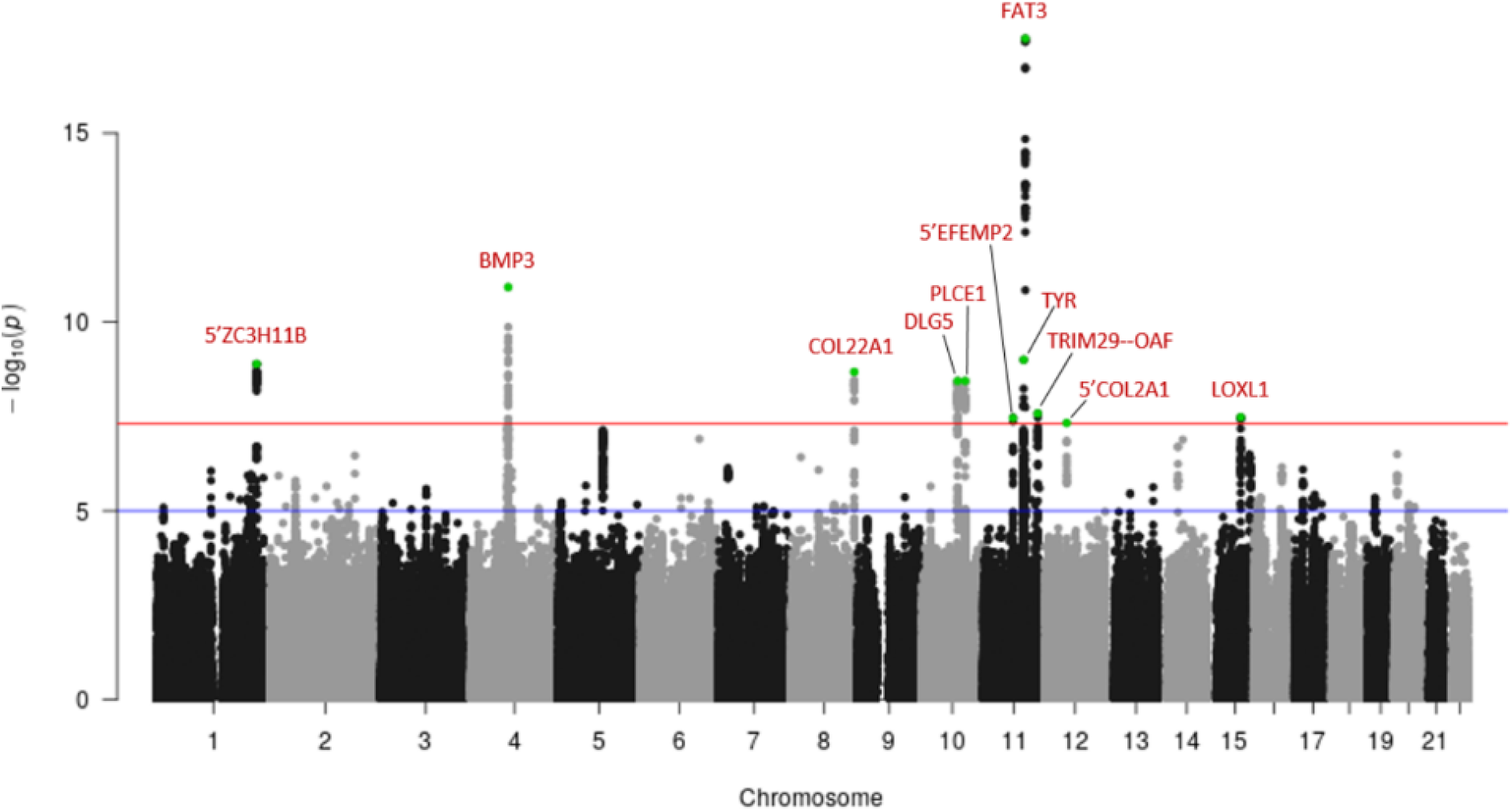
Manhattan plot following meta-analysis of genome-wide association studies using UK Biobank cases (RD-SR-ICD, N=3977) and clinically ascertained RRD cases (N=2164). The y axis represents the negative logarithm (base 10) of the meta-analysis P-values and the x axis the location of variants tested. Loci displaying association reaching the genome-wide significant threshold (red line, P < 5 × 10^−8^) are indicated in red, with index variant (lowest P-value within a locus) indicated in green. Suggestive level of significance (P < 10^−5^) is indicated by a blue line.

Gene-set analysis, with the genes prioritised from variants or genes associated at genome-wide significant level, highlights significant enrichment for genes encoding proteins with “receptor serine threonine kinase binding function” (*BMP2* and *3*) and “basement membrane components” (*EFEMP2*, *COL2A1* and *LOXL1*). Both of these molecular signature pathways are strengthened when more FUMA prioritised genes, corresponding to the lower GWAS P-value threshold of 10^−5^, are considered (S12 Table). The former extends to BMP2, 3 and 4 (P_enrichment_ =1.51 10^−3^) and the latter to *EFEMP2*, *COL2A1*, *LOXL1*, *EFEMP1*, *FRAS1*, *LAMA2* and *ACHE*, becoming the most statistically supported (P_enrichment_ =2.77×10^−4^) and unambiguous (all genes are on different chromosomes) molecular pathway. A trait-GWAS enrichment performed by FUMA (GWAS catalogue release r2018-02-06) reinforces evidence of overlap with myopia with six prioritised genes from the RD meta-analysis amongst the 29 genes annotated to a refractive error GWAS [42]: *GRIA4*, *RBFOX1*, *BMP2*, *BMP3*, *LAMA2*, *CHD7* (P_enrichment_ =4.66 10^−7^), and six additional RD genes present in other myopia-related traits-GWAS gene sets (S13 Table). Four genes (*DUSP12*, *BACS3*, *EFEMP1* and *PLCE1*) were in addition flagged as part of glaucoma endophenotypes GWAS gene-sets.

Replication was sought for the index variants of the eleven loci reaching genome-wide significant level of association by performing association analysis in the large 23andMe dataset. In this dataset, 9171 participants had self-reported retinal detachment, according to questionnaire administrated on a web based interface. Six associations, at *ZC3H11B*, *BMP3*, *COL22A1*, *PLCE1*, *TYR* and *FAT3* replicate at a Bonferroni corrected significance threshold of 0.0045 (P.val=0.05/11) (Table 3). In all the eleven look up, the alleles associated with increased retinal detachment risk were the same as in the GWAMA.

**Table 3.** Replication of retinal detachment associations in the 23andMe dataset

| SNP | Locus | EA | ALT | Effect | SE | P |
| --- | --- | --- | --- | --- | --- | --- |
| 1: 219759682<br>rs4373767 | **5' ZC3H11B** | T | C | 0.0444 | 0.0154 | **4.03x10<sup>-3</sup>** |
| 4:81952637<br>rs74764079 | **BMP3** | T | A | -0.1923 | 0.0450 | **3.02x10<sup>-5</sup>** |
| 8:139909653<br>rs11992725 | **COL22A1** | G | A | 0.1045 | 0.0160 | **7.05x10<sup>-11</sup>** |
| 10:79579222<br>rs1248634 | DLG5 | G | A | -0.0263 | 0.0170 | 0.122 |
| 10:96038686<br>rs11187838 | **PLCE1** | G | A | -0.0622 | 0.0152 | **4.51x10<sup>-5</sup>** |
| 11:65640906<br>rs7940691 | 5' EFEMP2 | T | C | 0.0047 | 0.0160 | 0.766 |
| 11:88911696<br>rs1042602 | **TYR** | C | A | -0.0514 | 0.0158 | **1.17x10<sup>-3</sup>** |
| 11:92601909<br>rs10765567 | **FAT3** | T | A | -0.1351 | 0.0160 | **1.64x10<sup>-17</sup>** |
| 11:120030227<br>rs11217712 | TRIM29---OAF | T | G | -0.0302 | 0.0165 | 6.73x10<sup>-2</sup> |
| 12:48404636<br>rs9651980 | 5' COL2A1 | T | C | 0.0508 | 0.0272 | 6.01x10<sup>-2</sup> |
| 15:74241624<br>rs4243042 | LOXL1 | T | A | -0.0098 | 0.0152 | 0.519 |

EA is the allele on the plus strand for which the effect is reported, ALT the alternate allele. Associations were tested using logistic regression, therefore effect are log of the odds ratio for retinal detachment. In bold, significant replication.

## Discussion

Combining the UK Biobank dataset with two independent datasets of clinically ascertained cases allowed a meta-analysis to be performed with close to 6000 retinal detachment cases, a case number required to power genetic analysis of complex traits, for which effect sizes are typically small[43]. A narrower definition of disease as in the clinically ascertained samples may provide better effect size estimates, but given that the largest collections of cases worldwide, to our knowledge, are the ones we have assembled and are still of modest size, power is very limited to identify robust statistical evidence for association using these collections alone. Indeed, the present study suggests a “winners curse” for several of the previously reported RRD associations.

We showed that the self-reported cases of retinal detachment in UK Biobank displayed characteristic epidemiologic features well-established for RRD. There is growing evidence that self-reported disease information can help increase power of genome-wide association studies [28,44]. For example, there was a remarkable similarity of findings between the 23andMe analysis using a simple online questionnaire for refractive errors and the CREAM consortium analysis using detailed measurements [44]. Data from the hospitalisation codes is not devoid of imprecision. In UK Biobank, over 25% of the RD cases were coded as H33.2, which represents serous retinal detachment. This is a far greater percentage than clinical experience would suggest. Although epidemiological data is available for RRD [1] no such data exists for serous RD due to the extreme rarity of its presentation. Furthermore, seven individuals recruited for the Scottish RRD study were also volunteers for the Scottish population-based GS:SFHS study. These individuals were definitively phenotyped as RRD by consultant surgeons, but they were all allocated H33.2 by the hospitalisation codes. This strongly suggests that a widespread systematic error has been made with regard to the specific code for subtypes of RD (H33.2 and H33.0), justifying our combination of these two codes for the purpose of analysis.

Despite the possibly increased heterogeneity in the retinal detachment phenotypes, the GWAS meta-analysis (GWAMA) yielded some consistent effects across sets and association with retinal detachment was replicated for six genome-wide significant index SNPs in the 23andMe dataset, where definition of cases was entirely self-reported. It is noteworthy that p-values in the replication were generally higher than in the GWAMA despite a larger number of cases: this could be due to a loosest phenotype definition, a web-based setting, younger median age of participants so that controls could be cases at older age with higher probability, or analysis method, a linear (logistic) regression method that may be less powerful in large sample size study than the mixed model setting[45].

Many of the implicated genes have function that appear very pertinent to retinal detachment aetiologies, validating the approach taken.

The locus near the well-established type 1 Stickler syndrome gene, *COL2A1,* was amongst the loci reaching genome-wide significance in the RD GWAMA, and index variant displayed a suggestive replication P-value of 0.06 in the 23andMe dataset. The mechanisms by which type II collagenopathy can favour RRD have been convincingly documented [6,46]. The 5’*COL2A1* locus has hallmarks of regulatory potential (open chromatin state observed in many cell lines-S8 Table) but none of the catalogued eQTLs in this region have been associated with the COL2A1 gene (S11 Table), showing the difficulty and limitations (e.g no eye tissue represented in eQTL databases) of functional interpretation and establishment of definite target gene(s) without follow-up experiments. Of note, patients with Stickler syndrome, at high risk of retinal detachment, are frequently myopic and commonly develop cataracts. To explore how the well-recognized epidemiological intertwining of these conditions was reflected in the genetic findings, we carried out GWAS on high myopia and on cataract operations in the UK Biobank dataset, to complement the available results from well powered GWAS available for common myopia or refractive error [33,47,48].

There was a clear myopia-related genetic load in line with expectation. Risk of chorio-retinal abnormalities increases with the severity of myopia and greater axial length [49,50,51]. This is a cause for concern, as myopia and, concomitantly, high myopia are showing significant increases in global prevalence [52]. Of note, not all loci strongly associated with high myopia confer increased RD risk, and those relating to increased elongation of the eye and basal membrane remodelling likely to be most relevant to RD. One of the six replicated RD locus, 5’ of *ZC3H11B*, is a newly identified refractive error locus[36] and has been associated with ocular axial length[53] and high myopia, in the Han Chinese population[54]. The genetic overlaps that large studies allow will increasingly inform on mechanisms for subsets of genes implicated, which are likely to act through diverse pathways, and may ultimately help differentiate myopic individuals at higher risk for developing specific pathological and potentially blinding complications.

The epidemiological associations of high myopia with cataract incidence and of cataract operations with RD were also well reflected in the high genetic overlap between all these conditions measured by the pairwise genetic correlations.

In a Dutch study [20], familial aggregation data had suggested that RRD genetic risk must extend beyond that contributing to myopia risk. In agreement, three of the replicated RD loci, *FAT3, TYR* and *COL22A1*, did not appear to exert their effect on RD primarily through myopia or cataract. *FAT3* and *COL22A1* were specifically associated with retinal detachment following look up of published data or of UK Biobank phenotypes GWAS repositories. FAT3 encodes for an atypical cadherin. Its pattern of expression is wide but its role in the retina is well established in mice. Mice homozygous for a knock-out allele exhibit abnormal amacrine cells displaying non polarised dendritic extensions, creating an additional synaptic layer in the retina [55]. It is possible that this altered structure would make the retina more prone to breakage, in keeping with the suggestion from the GWAS sensitivity analyses based on hospital records sub-codes that *FAT3* may be concerned with the retinal breaks.

Despite likely misclassifying of most H33.2 codes as discussed above, it is possible that the *COL22A1* association is stronger with serous detachments. *COL22A1* encodes for the collagen, type XXII, alpha 1 chain, a basement membrane collagen with a very restricted pattern of tissue expression [56]. It is highly expressed in the retina (mouse and human tissues) where its function is unknown. Its role in myotendinous junction (MTJ) has been well explored in zebrafish where knockdown of COLXXII expression resulted in contraction-induced muscle fiber detachment [57]. MTJ are rich in extra cellular matrix components and interaction of COLXXII collagen to integrin suggested in MTJ [57] points to a role in cell-ECM adhesion.

The TYR association is likely to act through variation of pigmentation in the retinal pigmented epithelium (RPE), the single cell layer of polarised cells that plays a crucial role in maintenance of the neural retina. Reported associations of the RD-associated variant with eye colour is less strong than that with skin pigmentation and variants with stronger effects on eye colour are not/or less associated with RD. While a large epidemiological study on prevalence of RRD in albino patients, whose tyrosinase is completely inactive, is unavailable, patho-physiologies that may concur with retinal detachment have been discussed[58], including increased posterior vitreous detachment incidence following increased eye-movement. Characterisation of an allelic series at the Tyr locus in mice suggests that albino adult mice retina are prone to detach in displayed sections compared to wild type mice[59]. Abnormal morphology of the RPE cells and impaired communication between the RPE and neuronal cells, affecting retinal ganglion cell (RGC) development, have been recently documented in albino mice embryos[60]. Compromised gap junctions between RPE and neural retina represents a compelling cause for retinal detachment risk.

The *PLCE1* locus had been flagged as the glaucoma endophenotype cup to disk ratio locus (index variants with RD index variant r2 0.6 D’=1)[61], and recently associated with primary open glaucoma[62] (index variant in complete LD with RD locus index variant), but has not been associated with IOP[63,64](the most powered of all glaucoma related GWAS). The *PLCE1* gene encodes for the phospholipase C epsilon 1 very strongly expressed in the retina where it has been suggested to regulate neuronal intracellular calcium levels and hence growth and differentiation signalling cascades[65]. Although glaucoma can develop secondary to a retinal detachment operation, epidemiology supports an increased prevalence of RD in glaucoma patients[29]. Our data supports the existence of genetic variations affecting both conditions.

The partitioning of variants based on their effects across traits is a very powerful tool to better understand and separate disease pathological pathways contributing to their aetiologies. For this approach to be thorough, it is clear that well powered studies are needed and for retinal detachment expanding the size of the study will be fruitful. With our study results largely driven by the availability of linked hospital records in UK Biobank, future expansion making use of new incident cases over time, the increased health records linkage, and the availability of similar datasets worldwide is a very promising way forward.

## Subjects and Methods

**Ethics statement:** All cohorts studied were approved by Research Ethics Committees as detailed below and all participants gave written informed consent. The UK Biobank study was conducted with the approval of the North-West Research Ethics Committee (Reference: 06/MRE08/65). Generation Scotland Scottish Family Health Study (GS:SFHS) has Research Tissue Bank status from the Tayside Committee on Medical Research Ethics (REC Reference: 15/ES/0040). The Scottish rhegmatogenous retinal detachment (RRD) study was approved by the Research Ethics Committee (MREC-06/MRE00/19) and the Moorfields RRD collection by Research Ethics Committee (10/H0703/97).

### Evaluation of retinal detachment self-report in UK Biobank

The criteria for self-reported retinal detachment (SR_RD) cases and controls based on the UK Biobank questionnaires are detailed in S2 Text. Proportions for conditions well documented to be associated with rhegmatogenous retinal detachment (RRD) were compared in cases and controls using a two-sided two-proportion z-test. Overlap with RD cases (ICD_RD) defined by relevant international classification of disease (ICD) codes from UK National Health Service (NHS) hospital patient records transcripts were examined. ICD codes relating to retinal detachment were ICD9 code 361 “retinal detachments and defects” and ICD10 codes H33 “retinal detachments and breaks”. In total 4777 (N=3913 ICD_RD and N=864 SR_RD with no available hospital record of detachment) were identified.

### Case-control datasets for genome-wide association analysis (GWAS)

#### UK Biobank sets

All analyses were restricted to the large subset of participants with genotype data passing quality control and indicative of white-British ancestry (S4 Text).

#### Retinal detachment

In the primary analysis, participants with retinal detachment ascertained from self report or linkage to hospital records were combined (RD-SR_ICD) as cases. Controls were participants who had declared not to have any contraindication for undergoing spirometry (as used in the evaluation of retinal detachment self-report), were not cases, and had not self-reported a history of retinal operation/vitrectomy. The discovery GWAS comprised N=3977 cases and N=360,233 controls; 10,000 controls recruited in the London centres of Barts, Croydon and Hounslow were not included in this analysis and used as independent controls in the analysis of the clinically ascertained RRD cases from the Moorfield collection.

The case:control ratio for the discovery retinal detachment analysis was therefore 1:91. GWAS was additionally performed using a more balanced 1: 3 case/control ratio for comparison, with controls matched to cases on gender and assessment centre. As effect sizes and P-values are influenced by the case/control ratio, a 1:10 ratio was also analysed for the look-up of UK Biobank top results in the clinically ascertained RRD cohorts results and their meta-analysis so that identical case/control ratio were compared or pooled together.

#### High myopia

N= 2737 high myopia cases (spherical equivalent refraction (SER) greater than −6 Diopters) and N=47635 controls (case:control ratio 1:17.4-Phenotype selection S2 Text), were contrasted.

#### Cataract operation

N= 21,679 cases and N=387,283 controls (Phenotype selection S2 Text-case:control ratio 1:17.9) were analysed.

### Clinically ascertained RRD cases and population-based controls

#### Scottish set

980 cases from the Scottish RRD study and 9,705 controls with no record linkage to a retinal detachment operation from GS:SFHS (SI Text-case:control ratio∼1:10) were analysed. 1,136,421 variants with high imputation quality (INFO >= 0.8) and MAF > 1% following genotyping analyses detailed in the S4 and S5 Text were tested.

#### London set

1184 cases recruited in London Moorfields eye hospital (S1 Text) and 10000 gender matched London centres recruited UK Biobank controls, defined as per UK Biobank analysis of RD, were analysed. In total, 4,727,220 variants with high imputation quality (INFO ≥ 0.9) and MAF > 1% following genotyping analyses detailed in the S4 and S5 Text were tested.

### Genome-wide association testing

Single variant GWAS was performed by testing for an additive effect at each reference allele within a linear mixed model to account for population relatedness and geographic structure. All analyses apart from one were performed using BOLT_LMM v1.3 [45,66] which implements fast parallelised algorithms to analyse large (N > 5000) datasets. The London RRD set analysis was run using GCTA[67] as low heritability estimation prevented BOLT_LMM v1.3 from carrying out the genome scan. Polygenic contribution to trait can be modelled using a mixture of Gaussians priors on SNP effect sizes (default model allowing uneven effect sizes) or a single-Gaussian prior (standard infinitesimal model) in BOLT_LMM v1.3. Reported association p-values are those obtained for the default BOLT_LMM model but all were identical or very close to those obtained under the standard infinitesimal model in our analyses. Covariates in the models were age (coded as 5 consecutive year of birth bins) and sex, and in addition, for the UK Biobank dataset, recruitment centre, genotyping array, genotyping batch and the 10 first components of the centrally performed UK Biobank principal components analysis. Only common to low frequency, MAF > 1%, variants, were analysed given the small size of the case samples and the risk of false positive associations for rarer variants in the unbalanced case:control samples analysed. In the UK Biobank dataset, recent simulations from the BOLT-LMM developers [45] show no inflation of type 1 error, for a significance threshold α of 5 10^−8^, at this MAF threshold with case fraction of 1% or higher.

Individual RD studies (each using a 1:10 case control ratio to harmonise effect sizes) summary statistics were meta-analysed using the inverse variance fixed effect scheme implemented in the software METAL[68]. Variants with Cochran’s Q-test heterogeneity of effect greater than 75% were filtered out.

### Replication

Replication analysis of 11 loci was performed using self-reported data from a GWAS including 9,171 cases and 406,168 controls of European ancestry, filtered to remove close relatives, from the 23andMe, Inc. customer database. All individuals included in the analyses provided informed consent and answered surveys online according to the 23andMe human subject protocol, which was reviewed and approved by Ethical & Independent Review Services, a private institutional review board (http://www.eandireview.com). Cases were defined as those who reported either retinal tears or retinal detachment; controls were defined as those who reported not having either retinal tears or retinal detachment.

All 11 variants assessed for replication were imputed in this study and were among the 21,747,472 variants passing quality control in the GWAS. Association test results were computed using logistic regression assuming additive allelic effects, including age, sex, the top five principal components of ancestry, and genotyping platform as covariates in the model.

### GWAS evaluation

Manhattan plots were drawn using the qqman library in R, and QQ plots using a local function. Regional association plots were plotted using LocusZoom: http://locuszoom.sph.umich.edu/ Inflation of type 1 error not accounted for by polygenicity was estimated using the LD score regression intercept [69] computed by the LDSC software. Z-scores calculated from the GWAS summary statistics from genotyped and well imputed (INFO >=0.8) variants were regressed on pre-calculated LD scores (of European ancestry from the 1000G phase 1 reference panel). For one GWAS, the Scottish RRD set, the ratio (*intercept* − 1)/(*mean*(*χ*^2^) − 1), which measures the proportion of the inflation in the mean *χ*^2^ that the LD Score regression intercept ascribes to causes other than polygenic heritability was greater than 20% and the LD score regression intercept was used as inflation factor to apply genomic control on the results[70].

A locus was defined as a region with significant variants within a 500kb region centred on the lead variant (that of the lowest P value in the region) and contiguous loci merged. Adjacent “loci” with variants displaying LD measure r^2^ > 0.2 across loci, e.g HLA region for cataract GWAS, were merged into one locus. Regions are flagged in tables when encompassing different blocks of variants with very different MAF (and hence low LD as measured by r2-https://www.ensembl.org/Homo_sapiens/Tools/LD/ using the 1000G phase_3:GBR dataset).

### Myopia genetic risk score

A myopia genetic risk score (GRS) was derived from 71 out of 140 genome-wide significant index variants for refractive error (P ≤ 5×10^−8^) from a large-scale meta-analysis of studies (CREAM and 23andMe meta-analysis) [36] independent from our discovery studies. The 71 variants passed QC in our three investigated cohorts. The GRS was constructed as the sum of the additive imputed dosage of the alleles increasing myopic refractive error, weighted by their effect on refractive error. Effect on retinal detachment risk was assessed using logistic regression with age and sex as covariates. The analysis using UK Biobank participants was restricted to those unrelated within the white-British ancestry subset (pairwise kinship coefficients lower than 0.0313).

### Functional annotation and gene mapping

Functional annotation was performed using FUMA [40] v1.3.1, a recently developed integrative tool using information from 18 biological data repositories. FUMA defines independent significant SNPS within loci if they are in low LD r2< 0.1 (using the 1000G Phase3 EUR reference panel [71]). These variants, as well as those, tested or not, in high LD (r^2^ >=0.8) with them in the 1000G EUR reference panel, were annotated for predicted protein-altering or regulatory features following annotations from ANNOVAR (http://annovar.openbioinformatics.org), RegulomeDB (http://www.regulomedb.org/), eQTL repositories (with note that none is available as yet for eye tissues) and the ChromHMM predicted 15 core chromatin states derived from ENCODE and ROADMAP repositories. SNP-trait known associations from the GWAS catalogue (r2018-02-06) were also part of FUMA annotations. The SNP2GENE function in FUMA identified putative target genes by physical mapping to the defined loci, or based on genes whose expression has been significantly associated with the considered GWAS variants or implicated by chromatin interaction data. eQTL repositories used were: GTEx v6 and v7, BIOS QTL browser and BrainSpan.

### Gene based analysis and Gene-set analysis

These analyses were performed using MAGMA [72] within FUMA. Gene-based P values are derived from an approximation of the sampling distribution of the mean of the x^2^ statistic for the SNPs in a gene, with gene coordinates that from NCBI 37.3 and LD pattern to account for dependency of SNPs P values that of the 1000G phase3 European reference data as input parameters. Genome-wide threshold of significance for the gene-based p-values was set to 2.74 ×10^−6^, following Bonferroni correction for the N=18246 genes evaluated.

Gene-based P-values converted to Z values and a between-gene correlation matrix are used as input to perform gene sets enrichment tests. [72]. These tests are based on expectation of an hypergeometric distribution for the null hypothesis of no enrichment. The raw p-values are adjusted following the Benjamini and Hochberg procedure (FDR set at 5%) to account for multiple testing within each gene-set categorie. Predefined gene sets from the molecular signatures database MsigDB v6.1 (http://software.broadinstitute.org/gsea/msigdb) were used.

### Traits Heritability and genetic correlations

The heritability attributable to the joint contribution of genetic variants tagged by the common variants tested, 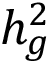, can be calculated using the variance components estimates from the models fitted to perform the GWAS. It was converted to a case:control ratio-independent heritability, on the scale of liability, 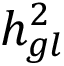 using the formula (1) discussed in[73] or (2) when controls were also selected using a trait threshold[74] for the analysis of high myopia)

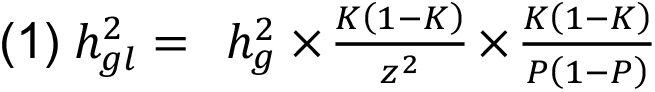

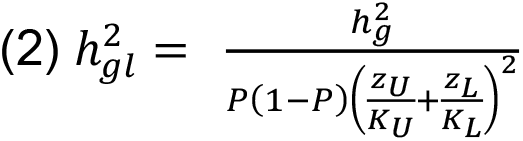

where P is the prevalence of the binary trait in the analysed sample (e.g for the largest retinal detachment analysis in UK Biobank: 3977/(360233+3977)=1.1%), K the prevalence in the population of matching age group as that of the UK Biobank population. We made the assumption that conditions’ prevalences in the UK Biobank sample are similar to that of the UK population; for retinal detachment, assuming K=P was close to a previous UK RRD estimate of incidence of 12 per 100,000 population-year[1] with median age of 59 year old in the sample. For high myopia, cases corresponded to the 3.76% upper tail of the refractive error distribution and controls to the 65.5% lower tail. z^2^ is the squared ordinate of a standard normal curve at the K quantile.

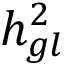 and 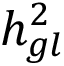 were also estimated using the linkage-disequilibrium (LD) score regression method[69]. Genetic correlations (r_g_) were those obtained by regressing the product of test statistics against LD-score[35] as implemented in the LDSC software.

### GWAS look-up

Four GWAS results repositories were used: the GWAS catalogue (r2018-02-06), PhenoScanner (as of March 2018) the Global Biobank Engine, Stanford, CA (URL: http://gbe.stanford.edu) (GBE) [March 2018] and the GeneATLAS, University of Edinburgh (URL: http://geneatlas.roslin.ed.ac.uk/) [March 2018]. In the Global Biobank Engine, over 2000 UK Biobank traits (including retinal detachment defined as in our discovery analysis) have been analysed in a first broad brush pass, on 337,000 unrelated participants of white British ancestry. The analysed traits include traits measured in smaller subsets of individuals in recall assessments, including the ophthalmologic assessment (e.g refractive error as a quantitative trait is analysed for N=82,752 participants). The GeneATLAS shows GWAS results for 778 traits measured in the complete set of UK Biobank participants of white British ancestry, using both related and unrelated individuals; results for the genotyped variants can be queried using the web interface.

## Acknowledgments

This research has been conducted using the UK Biobank Resource under application n^0^ 37502 formely 8842. We specially thank Mrs Lorraine Gillions for her help in the resource access process.

The work in this paper uses data provided by patients and collected by the NHS as part of their care and support.

Genotyping of the GS:SFHS DNA and Scottish RRD samples was carried out by the Genetics Core Laboratory at the Edinburgh Clinical Research Facility, University of Edinburgh. Genotyping of ∼1200 RRD participants recruited in Moorfields was performed at the Human Genomics Facility (HuGe-F) at Erasmus MC (www.glimdna.org). We are grateful to Prof André Uitterlinden, Dr Joyce van Meurs, Dr Fernando Rivadeneira, Dr Linda Broer and Mila Jhamai for coordinating the GSA genotyping effort. The authors would like to thank the Rivas lab for making the Global Biobank Engine resource available and the Tenesa group for the GeneATLAS. In addition, we thank the research participants of 23andMe for their contribution to this study and the following members of the 23andMe Research Team: Michelle Agee, Babak Alipanahi, Adam Auton, Robert K. Bell, Katarzyna Bryc, Sarah L. Elson, Pierre Fontanillas, Nicholas A. Furlotte, Barry Hicks, Karen E. Huber, Ethan M. Jewett, Yunxuan Jiang, Aaron Kleinman, Keng-Han Lin, Nadia K. Litterman, Matthew H. McIntyre, Kimberly F. McManus, Joanna L. Mountain, Elizabeth S. Noblin, Carrie A.M. Northover, Steven J. Pitts, G. David Poznik, J. Fah Sathirapongsasuti, Janie F. Shelton, Suyash Shringarpure, Chao Tian, Joyce Y. Tung, Vladimir Vacic, Xin Wang, Catherine H. Wilson. Finally, we would like to express our gratitude to Jonathan Marten for providing the plotting scripts for the genetic risk scores, Dr Amy Findlay for pointing out relevant past publications on albino mice and Dr Shona Kerr, “QTL in Health and Disease” programme project manager, for critical reading of the manuscript.

## Web Resources

**URLs**. UK Biobank Access Management System, http://www.ukbiobank.ac.uk/register-apply/; HRC variants, http://www.haplotype-reference-consortium.org/site; LocusZoom: http://locuszoom.sph.umich.edu/; NHGRI-EBI catalog of published genome-wide association studies, http://www.ebi.ac.uk/gwas/; Phenoscanner, http://www.phenoscanner.medschl.cam.ac.uk/phenoscanner; LDSC, https://github.com/bulik/ldsc; BOLT, https://data.broadinstitute.org/alkesgroup/BOLT-LMM/; FUMA, http://fuma.ctglab.nl/; Haploreg, http://archive.broadinstitute.org/mammals/haploreg/haploreg.php; LD calculation tool, https://www.ensembl.org/Homo_sapiens/Tools/LD/, CADD pre-calculated scores, http://cadd.gs.washington.edu/

## Supporting Information

**S1 Text**. Study populations

**S2 Text**. Phenotype selection

**S3 Text**. UK Biobank Eye and Vision Consortium Membership

**S4 Text**. Genotype preparation

**S5 Text**. Imputation to the Haplotype Reference Consortium panel for the clinical case-control dataset

**S1 Fig**: Study design flow chart.

**S2 Fig**: Manhattan and quantile-quantile plots for the UK Biobank retinal detachment primary GWAS (Self report and ICD10 H33 codes or ICD9 316 code; N=3977 cases of white British ancestry; N=360233 controls) using genotypes imputed to the Haplotype Reference Consortium reference.

**S3 Fig**. Manhattan plot of the retinal detachment GWAS in UK Biobank (Self report and ICD10 H33 codes or ICD9 316 code; N=3977 cases of white British ancestry) when a 1:3 case to recruitment-centre and sex-matched control ratio is used.

Significance thresholds corresponding to P-value=5 × 10^−8^ and 10^−5^ are represented respectively in red and blue.

**S4 Fig**. Manhattan plots of GWAS for conditions epidemiologically associated with retinal detachment using the UK Biobank data. Significance thresholds corresponding to P-values of 5 × 10^−8^ and 10^−5^ are represented respectively by red and blue lines.

**S5 Fig**. Regional association plots around the RD-SR-ICD UK Biobank loci in epidemiologically related traits

**S6 Fig**. Distributional properties of myopia genetic risk score and retinal detachment status

**S7 Fig**. Forest plot for the meta-analysis of effects on retinal detachment for the TYR missense variant rs1042602 (effect reported for the C allele). TE effect size in case control sets with a 1:10 case to control ratio

**S8 Fig**. Plots of the first two principal components in multidimensional data reduction of the genotypes available in both clinical cases and controls. The set of variants used for analysis were pruned to remove variants in high linkage disequilibrium. A. Scottish Study RRD cases (N=981) and GS:SFHS controls (N=9754) and B. Moorfield recruited RRD cases and UK Biobank participants recruited in London and defined as of white british ancestry by the UK Biobank central analysis team used as controls. Black circles represent control and red circles cases.

**S1 Table** Regions associated with retinal detachment in UK Biobank (RD-SR-ICD) using a P-value threshold of 5×10^−6^.

**S2 Table.** Regions associated with ICD10 code H33.0 in UK Biobank (N=1380 cases) using a P-value cut-off of 10^−6^

**S3 Table**. Genome-wide significant signals for high myopia in UK

**S4 Table**. Genome-wide significant signals for cataract operation in UK Biobank

**S5 Table**. Association look-up for index variants associated with RD-SR-ICD at a P-value cut-off of 5×10^−6^

**S6 Table**. Regions associated with RRD following meta-analysis of two clinically ascertained cases datasets, using a P-value threshold of 5×10^−6^.

**S7 Table**. Regions associated with retinal detachment in the combined UK Biobank and clinical datasets using a P-value threshold of 10^−6^.

**S8 Table**. Notable functional annotations for the lead and tagged variants underlying the 11 identified risk loci for retinal detachment (N=564 variants). Tagged variants in LD with lead variants (r^2^ >= 0.8) included tested as well as untested variants from the 1000G reference phase 3 within a 250 kb distance of lead variants. Variants with the strongest indications of functionality following annotation by FUMA are shown: with high CADD scores (CADD) which measure how deleterious a variant is predicted to be, with low RegulomeDB scores (RDB – which ranges from 1a for an eQTL variant with evidence of a transcription factor (TF) binding a motif for that TF and a DNase footprint to 7 for variant with no regulatory evidences) or/and with a 15 core chromatin states across 127 tissue/cell types from ENCODE and ROADMAP repositories (minChrState) score lower than 8 which indicates an open chromatin state. Nearest gene (nearestgene) and distance to it (Dist) are based on ANNOVAR annotation using Ensembl gene 85 (GRCh37 human genome assembly). Whether the variant is or not within the physical boundary of the locus is indicated by the indicator posMapFilt (0 for no 1 for yes) and whether it is an eQTL by the indicator poseqtlFilt. eQTL evidences were from GTEx v6, GTEx v7, BIOS QTL and BrainSpan.

**S9 Table**. Reported trait associations for the lead variants underlying the 11 identified risk loci for retinal detachment. Four GWAS results repositories were used: the GWAS catalogue (r2018-02-06), PhenoScanner (as of March 2018) the Global Biobank Engine (URL: http://gbe.stanford.edu) (GBE) [March 2018] and the GeneATLAS (URL: http://geneatlas.roslin.ed.ac.uk/) [March 2018]. Significance: P-value (P) threshold * P < 0.05, ** P < 1×10^−5^, *** P< 5×10^−8^; Categorie: 1 = ocular with no known RD association, 2 = ocular with known retinal detachment risk,3 = retinal detachment, 0 otherwise. For each resource, any phenotype associated at the standard genome-wide significant P-value (P < 5×10^−8^) is listed. If no such trait is reported, the top associated trait is reported, and any ocular trait nominally associated is listed.

**S10 Table**. Suggestive gene-based associations (P < 10^−5^) performed using MAGMA and the RD meta-analysis summary statistics. Novel RD genes with P-value exceeding the genome-wide significant threshold (2.74×10^−6^ following Bonferroni correction for the N= 18246 genes tested) are highlighted in bold. Positions (start and stop) follow GRCh37 human genome assembly.

**S11 Table**. Annotations for the candidate genes selected at each identified RD-associated genetic locus by FUMA (SNP2GENE) or by the gene-based association analysis. SNP2GENE identified genes either by physical mapping to the defined loci or based on eQTL evidences linking an associated variant within the locus to the gene. The maximum CADD score for variants mapping to each gene (posMapMaxCADD), the strength of the eQTL association (eqtlMapminQ) and source of the eQTL studies (eqtlMapts) are indicated. The genes in bold are physically located in the loci. Transcript levels in human ocular tissue are from the Eye Integration v0.62 database where processed deep RNA sequencing data from three ocular tissues are deposited and the Ocular Tissue database (https://genome.uiowa.edu/otdb/) where transcription levels in a wider range of ocular tissues has been assessed using a chip-array. Mouse mutant phenotypes were looked up in the Mouse Genome Informatics resource (http://www.informatics.jax.org/) and the International Mouse Phenotyping Consortium (http://www.mousephenotype.org/) as of March 2018 and human associated conditions in OMIM (https://www.omim.org/) with HPO annotation of commonly associated features from https://rarediseases.info.nih.gov/

**S12 Table**. Top (adjP < 0.01) gene-set or pathway enriched using MAGMA on the RD meta-analysis results. The prioritized genes from the meta-analysis genome-wide significant variants or genes (GWset) and by all variants associated at a lower P-value threshold, 10^−5^, (SUGset) were used as input. The GWset includes the 49 genes identified by FUMA SNP2GENE and the 4 additional independent genes detected by gene-based analysis. The SUGset includes 283 genes. GO:Gene ontology; cc: cellular component; mf: molecular functions; bp: biological processes; N:number of genes in gene set.* denotes physically close genes within gene set hence likely biased analysis

**S13 Table**. Enriched GWAS-based gene sets in the RD meta-analysis results. RD associated variants with a P-value below the threshold of 10^−5^ were used as input for FUMA annotation. GWAS for ocular traits are highlighted in bold

## Data reporting

GWAS statistics for all analyses presented here are available from the University of Edinburgh Digital Repository DataShare: (DOI in progress)

