## supporting information for "Insight into the genetic aetiology of retinal detachment by combining small clinical and large population-based datasets"

### S1 Text: Study Populations

All cohorts studied were approved by Research Ethics Committees as detailed below and all participants gave written informed consent.

#### The UK Biobank (UKBB)

The UK Biobank is a large-scale prospective study established by the Medical Research Council, Department of Health, Wellcome Trust medical charity, Scottish Government and Northwest Regional Development Agency[1]. Between 2006 and 2010, close to 500,000 people (273,467 female/229,175 male) were recruited, and genome-wide genotype data on the full set has recently been made available to the international community [2]. A detailed description of the study and its access process are available online (http://www.ukbiobank.ac.uk/resources/). A baseline questionnaire was undertaken in the 22 assessment centres across the UK. All UK residents aged 40 to 69 years who were registered with the National Health Service (NHS) and living up to 25 miles from a study centre were invited to participate. The study was conducted with the approval of the North-West Research Ethics Committee (Reference: 06/MRE08/65). Phenotypes and genotypes used in the manuscript were obtained through approved application number 8842.

#### Generation Scotland Scottish Family Health Study (GS:SFHS)

GS:SFHS is a genotyped family-based cohort recruited from the general population across Scotland, UK. Over 24,000 adults, aged 18-98 years, were recruited from 2006 to 2011, with broad and enduring written informed consent for biomedical research [3]. GS:SFHS has Research Tissue Bank status from the Tayside Committee on Medical Research Ethics (REC Reference: 15/ES/0040). Specific consent was obtained at recruitment from 23,603 participants for GS:SFHS study data to be linked to their Scottish National Health Service (NHS) records, using their Community Health Index (CHI) number, allowing healthcare records for individuals to be linked across time and location[4]. Study description and access process are available online (www.generationscotland.org). The request to use GS:SFHS participants as controls was made through application GS13127.

**The Scottish rhegmatogenous retinal detachment (RRD) study**

The Scottish RRD study is a cross-sectional study that sought to recruit all incident cases of primary RRD presenting to each vitreo-retinal surgical site in Scotland between November 2007 and November 2009 [5,6]. 1,244 incident cases of primary RRD from 6 centres across Scotland were recruited, representing over 95% of all operated cases in Scotland during this time period. The diagnosis of RRD was made by vitreo-retinal consultant specialists based on a case definition of "a full thickness break in the neurosensory retina with a surrounding area of sub-retinal fluid extending greater than 2 disc diameters". Blunt traumatic cases of RRD were included, penetrating trauma was excluded. All other types of retinal detachment (exudative, tractional and combined) were excluded. DNA from 912 Scottish RRD cases have been genotyped using the Illumina CNV370v3-Quad genome-wide SNP array as previously described and analysed [7]. 121 samples, including four passing genotyping quality control in that primary analysis, were newly genotyped using the HumanOmniExpressExome-8 v1.2 BeadChip. 999 samples, overlapping with both of these datasets, had also been genotyped for an exome variants analysis using the Illumina HumanExome-12v1-1_A array. The study was approved by Research Ethics Committee (MREC-06/MRE00/19)

#### The Moorfields RRD collection

A case collection was established in the Moorfields Eye Hospital, London. Clinical diagnosis of RRD was confirmed by consultant vitreoretinal surgeons. A research nurse and clinical fellow recruited and collected a blood sample from all eligible and consenting incident cases of RRD presenting or referred to Moorfields Eye Hospital between August 2010 and December 2012, totalling 1996 participants. The blood samples were frozen on site, DNA extracted, barcoded and stored at monthly intervals in the KBioscience, (later LGC), facility. The criteria for cases identification were the same as used in the Scottish RRD study. 1,248 of the cases collected were genotyped for GWAS using the Illumina Global Screening Array (GSA) which encompasses close to 700,000 markers. The study was approved by Research Ethics Committee (10/H0703/97)

### S2 Text-Phenotype selection

#### UK Biobank

#### Self reported retinal detachment cases and controls

An indicator variable was created in the R software environment (https://www.r-project.org/) to score instances of code “1281= retinal detachment” extracted from data field f20002, encoding self-reported non cancer illnesses recorded during verbal interviews at UK Biobank assessment centres. The initial assessment visit at which participants were recruited was between 2006 and 2010; a subset had two subsequent assessment visits (2012-2013 and 2014+). Indicator variables for other self-reported conditions and operations associated with RRD were also created: T2D (code 1223),

T1D (code 1222), Diabetic eye disease (code 1276), glaucoma (code 1277), cataract (code 1278), eye trauma (code 1279), and amongst self-reported operations (field 20004) retinal operation/vitrectomy (code 1437) and cataract surgery (code 1435). Additionally, macula degeneration (f20002 code 1528) was extracted. Also gathered at the same time from most participants, "have ever had Eye problems other than wearing glasses" from field 2227 and "age started wearing glasses or contact lenses"(f2217) when reason for glasses/contact lenses (f6147) is " for short-sightedness"(code 1) were extracted.

Controls were chosen as participants unlikely to have retinal detachment based on a question given at recruitment about any contraindication for undergoing spirometry (data field f3088), one contraindication being "history of retinal detachment and surgery to eyes". Answers to that question were thus available for all participants (in our data release N=502,618) and are coded Yes, No, Unsure or NA (if missing). A categorical variable, combining the responses at the three assessment visits, was created with values NA if answer is missing at all of the three instances (N=1693), No for any combination of No and NA, Yes for any combination of Yes and NA, Yes Recent for Yes in the last non NA instance, Unsure for any combination of NA or Yes and Unsure, No Likely for all other combination. The definition of controls was set as individuals with value No (N=457 255) for this variable. Those controls were used in the assessment of self-report, and a subset used as controls in the RD GWAS analysis. For a small subset of 2,014 participants, the explicit reason for not undertaking spirometry is provided (f20042.0.0) and of those, 83 participants declared "retinal detachment/ eye problem" (code 508). However, those entries did not seem reliable and were not included as additional self-reported cases: 80 out of the 83 had answered no to have (/have had) a condition preventing spirometry in field3088 and only 9 individuals amongst the 83 (11%) self-reported a retinal operation/vitrectomy which would be expected with retinal detachment.

**Hospital records based retinal detachment cases**

Individuals were identified as cases if an operation was ever recorded with an ICD code relating to retinal detachment: ICD9 codes (UKBB fields f41203 and f41205) 361 “retinal detachments and defects” and ICD10 codes (f41202 and f41204) H33 “retinal detachments and breaks”. ICD10 code H33 encompasses subclasses H33.0 (retinal detachment with retinal break) which should corresponds to RRD, H33.1 (retinoschisis and retinal cysts), H33.2 (serous retinal detachment), H33.3 (retinal breaks without detachment) H33.4 (traction detachment of the retina) and H33.5 (other retinal detachments).

**High myopia**

Refractive error measured using a TomeyRC-500 device is available for 133,265 UK Biobank participants. High myopia cases were defined as of average (from both eyes) Spherical Equivalent Refraction (SER) lower than -6 Diopters, with SER derived as Spherical power (f5084 right eye, f5085 left eye) plus half Cylindrical power (f5087, f5086). Eyes for which the refractometry was unreliable (f5091, f5090) or affected by injury or trauma (f5419) were not included. Controls were individuals with average SER no lower than -0.5 Diopters and no history of cataract surgery (f5324), corneal graft surgery (f5328), or refractive laser surgery (f5325).

**Cataract operation**

Cases were defined as participants with a self-reported cataract operation (f20004 code 1435) or/and a hospital record including an ICD10 code H25 or H26. Controls were participants who were not cases.

### Scottish RRD cases and controls

All but one of the 981 Scottish RRD cases passing genotyping QC (Supplemental methods 1) were used in the analysis. One individual was removed as noted to have a systemic condition associated with RRD, Marfan Syndrome. Of the 9761 GS:SFHS controls who passed genotyping QC, seven cases as well as forty nine additional participants linked to an ICD10 code H33 were excluded, leaving 9705 controls for the analysis. Linkage to hospital inpatients admissions dated from December 2016 and include most of the GS:SFHS cohort. The seven cases who were also GS:SFHS participants (based on 100% genotype sharing) were all linked to an ICD-10 H33 code. Of note, all were linked to sub-code H33.2 (serous detachment) and only two additionally recorded with a H33.0 sub-code (retinal detachment with retinal break), the expected descriptor of RRD, at a later hospital visit.

### 23andMe

Cases and controls were selected based on participant responses to the two following questions: "Have you ever been diagnosed by a doctor with any of the following vision conditions/retinal detachment/retinal tears?" and "Have you ever been diagnosed with retinal detachment that was NOT caused by a physical trauma, such as a concussion or blow to the eye area?". Participants who answered "yes" to either question were defined as cases; participants who answered "no" to at least one of these questions and did not answer "yes" to either were defined as controls, otherwise they were classified as NA. The choice of answers was “Yes”, “No”, “I don’t know/I’m not sure”.

### S3 Text. UK Biobank Eye and Vision Consortium Membership

• Prof Tariq ASLAM - Manchester University, Manchester, UK
• Prof Sarah BARMAN - Kingston University, London, UK
• Prof Jenny BARRETT - University of Leeds, Yorkshire, UK
• Prof Paul BISHOP - Manchester University, Manchester, UK
• Mr Peter BLOWS - NIHR Biomedical Research Centre, Moorfields Eye Hospital NHS Foundation Trust and UCL Institute of Ophthalmology, London, UK
• Dr Catey BUNCE - King's College London, London, UK
• Dr Roxana CARARE - University of Southampton, Southampton, UK
• Prof Usha CHAKRAVARTHY - Queens University Belfast, Belfast, Ireland
• Miss Michelle CHAN - NIHR Biomedical Research Centre, Moorfields Eye Hospital NHS Foundation Trust and UCL Institute of Ophthalmology, London, UK
• Dr Sharon CHUA - NIHR Biomedical Research Centre, Moorfields Eye Hospital NHS Foundation Trust and UCL Institute of Ophthalmology, London, UK
• Prof David CRABB – City, University of London, London, UK
• Mrs Philippa CUMBERLAND - UCL Great Ormond Street Institute of Child Health, London, UK
• Dr Alexander DAY - NIHR Biomedical Research Centre, Moorfields Eye Hospital NHS Foundation Trust and UCL Institute of Ophthalmology, London, UK
• Miss Parul DESAI - NIHR Biomedical Research Centre, Moorfields Eye Hospital NHS Foundation Trust and UCL Institute of Ophthalmology, London, UK
• Prof Bal DHILLON - University of Edinburgh, Scotland, UK
• Prof Andrew DICK - University of Bristol, Bristol, UK
• Dr Cathy EGAN - NIHR Biomedical Research Centre, Moorfields Eye Hospital NHS Foundation Trust and UCL Institute of Ophthalmology, London, UK
• Prof Sarah ENNIS - University of Southampton, Southampton, UK
• Prof Paul FOSTER - NIHR Biomedical Research Centre, Moorfields Eye Hospital NHS Foundation Trust and UCL Institute of Ophthalmology, London, UK
• Dr Marcus FRUTTIGER - NIHR Biomedical Research Centre, Moorfields Eye Hospital NHS Foundation Trust and UCL Institute of Ophthalmology, London, UK
• Dr John GALLACHER - University of Oxford, Oxford, UK
• Prof David (Ted) GARWAY-HEATH - NIHR Biomedical Research Centre, Moorfields Eye Hospital NHS Foundation Trust and UCL Institute of Ophthalmology, London, UK
• Dr Jane GIBSON - University of Southampton, Southampton, UK
• Mr Dan GORE - NIHR Biomedical Research Centre, Moorfields Eye Hospital NHS Foundation Trust and UCL Institute of Ophthalmology, London, UK
• Prof Jeremy GUGGENHEIM - Cardiff University, Wales, UK
• Prof Chris HAMMOND - King's College London, London, UK
• Prof Alison HARDCASTLE - NIHR Biomedical Research Centre, Moorfields Eye Hospital NHS FoundationTrust and UCL Institute of Ophthalmology, London, UK
• Prof Simon HARDING - University of Liverpool, London, UK
• Dr Ruth HOGG - Queens University Belfast, Belfast, Ireland
• Dr Pirro HYSI - King's College London, London, UK
• Mr Pearse A KEANE - NIHR Biomedical Research Centre, Moorfields Eye Hospital NHS Foundation Trust and UCL Institute of Ophthalmology, London, UK
• Prof Sir Peng Tee KHAW - NIHR Biomedical Research Centre, Moorfields Eye Hospital NHS Foundation Trust and UCL Institute of Ophthalmology, London, UK
• Mr Anthony KHAWAJA - NIHR Biomedical Research Centre, Moorfields Eye Hospital NHS Foundation Trust and UCL Institute of Ophthalmology, London, UK
• Mr Gerassimos LASCARATOS - NIHR Biomedical Research Centre, Moorfields Eye Hospital NHS Foundation
Trust and UCL Institute of Ophthalmology, London, UK
• Prof Andrew LOTERY- University of Southampton, Southampton, UK

• Prof Phil LUTHERT - NIHR Biomedical Research Centre, Moorfields Eye Hospital NHS Foundation Trust and
UCL Institute of Ophthalmology, London, UK
• Dr Tom MACGILLIVRAY - University of Edinburgh, Scotland, UK
• Dr Sarah MACKIE - University of Leeds, Yorkshire, UK
• Prof Keith MARTIN - University of Cambridge, Cambridge, UK
• Ms Michelle MCGAUGHEY - Queen’s University Belfast, Belfast, Ireland
• Dr Bernadette MCGUINNESS - Queen’s University Belfast, Belfast, Ireland
• Dr Gareth MCKAY - Queen's University Belfast, Belfast, Ireland
• Mr Martin MCKIBBIN - Leeds Teaching Hospitals NHS Trust, Yorkshire, UK
• Dr Danny MITRY – NIHR Biomedical Research Centre, Moorfields Eye Hospital NHS Foundation Trust and UCL Institute of Ophthalmology, London, UK & Royal Free Hospital, London, UK
• Prof Tony MOORE - NIHR Biomedical Research Centre, Moorfields Eye Hospital NHS Foundation Trust and UCL Institute of Ophthalmology, London, UK
• Prof James MORGAN - Cardiff University, Wales, UK
• Ms Zaynah MUTHY – NIHR Biomedical Research Centre, Moorfields Eye Hospital NHS Foundation Trust and UCL Institute of Ophthalmology, London, UK
• Mr Eoin O'SULLIVAN - King's College Hospital NHS Foundation Trust, London, UK
• Dr Chris OWEN - St George's, University of London, London, UK
• Mr Praveen PATEL - NIHR Biomedical Research Centre, Moorfields Eye Hospital NHS Foundation Trust and UCL Institute of Ophthalmology, London, UK
• Mr Euan PATERSON - Queens University Belfast, Belfast, Ireland
• Dr Tunde PETO - Queen's University Belfast, Belfast, Ireland
• Dr Axel PETZOLD - UCL Institute of Neurology, London, UK
• Prof Jugnoo RAHI - UCL Great Ormond Street Institute of Child Health, London, UK
• Dr Alicja RUDNICKA - St George's, University of London, London, UK
• Mr Jay SELF - University of Southampton, Southampton, UK
• Prof Sobha SIVAPRASAD - NIHR Biomedical Research Centre, Moorfields Eye Hospital NHS Foundation Trust and UCL Institute of Ophthalmology, London, UK
• Mr David STEEL - Newcastle University, Newcastle, UK
• Mrs Irene STRATTON - Gloucestershire Hospitals NHS Foundation Trust
• Mr Nicholas STROUTHIDIS - NIHR Biomedical Research Centre, Moorfields Eye Hospital NHS Foundation Trust and UCL Institute of Ophthalmology, London, UK
• Prof Cathie SUDLOW - University of Edinburgh, Scotland, UK
• Dr Caroline THAUNG - NIHR Biomedical Research Centre, Moorfields Eye Hospital NHS Foundation Trust and UCL Institute of Ophthalmology, London, UK
• Miss Dhanes THOMAS - NIHR Biomedical Research Centre, Moorfields Eye Hospital NHS Foundation Trust and UCL Institute of Ophthalmology, London, UK
• Prof Emanuele TRUCCO - University of Dundee, Scotland, UK
• Prof Adnan TUFAIL - NIHR Biomedical Research Centre, Moorfields Eye Hospital NHS Foundation Trust and UCL Institute of Ophthalmology, London, UK
• Dr Veronique VITART - University of Edinburgh, Scotland, UK
• Prof Stephen VERNON – Nottingham University Hospitals NHS Trust, Nottingham, UK
• Mr Ananth VISWANATHAN - NIHR Biomedical Research Centre, Moorfields Eye Hospital NHS Foundation Trust and UCL Institute of Ophthalmology, London, UK
• Miss Cathy WILLIAMS - University of Bristol, Bristol, UK
• Dr Katie WILLIAMS - King's College London, London, UK
• Prof Jayne WOODSIDE - Queen's University Belfast, Belfast, Ireland
• Dr Max YATES - University of East Anglia, Norwich, UK
• Ms Jennifer YIP - University of Cambridge, Cambridge, UK
• Dr Yalin ZHENG - University of Liverpool, London, UK

### S4 Text. Genotype preparation

#### UK Biobank

The data release includes genotypes for 97.2% (N=488,377) of the UK Biobank participants. Genotyping was performed using two related Affymetrix arrays of close to 800,000 variants: the UK BiLEVE array for the first 50,000 samples and the Affymetrix Axiom® for the remaining samples (http://www.ukbiobank.ac.uk/wp-content/uploads/2014/04/UK-Biobank-Axiom-Array-Content-Summary-2014.pdf). Full description of the genotype data quality controls and subsequent imputation carried out using the HRC+UK10K panel can be found on http://www.ukbiobank.ac.uk/wp-content/uploads/2014/04/imputation_documentation_May2015.pdf. 968 samples have been flagged as failing the centrally performed quality controls for heterozygosity or/and missingness[2], leaving 487,409 individuals available for analysis. Here, we further excluded samples flagged as having a mismatch between self-reported and genotype-derived gender (N=378) or showing putative sex chromosome aneuploidy (N=652). We also excluded individuals of non-white-British ancestry (N=78674 in the genotype release) from our analyses. Following the identification of a problem with the non Haplotype Reference Consortium (HRC)[8] imputed variants in the July 2017 release of the data which we were using, those were filtered out from the analysis results and only HRC sites are presented for the imputed data analysis; the HRC site list is available at http://www.haplotype-reference-consortium.org/site.

#### Scottish RRD cohort

For each array, the genotype clusters for individuals with call rate > 90% were examined in genome studio using standard recommendations [9]. Genotype files in plink format, on TOP strand notation, were generated using the genome studio PLINK plugin v2.1.4. SNPs with a call rate less than 99%, or deviation from Hardy-Weinberg equilibrium exceeding p-value of 1 x $10^{-6}$ in significance were excluded. Individuals with a call rate less than 98% were removed, as well as samples identified as duplicates and those with mismatch between self-reported and genotype-derived gender. Variant naming was harmonised between arrays using positions on the same reference build of the genome (hg19 b37). Duplicate markers, mismatches of chromosome, position, strand or alleles were identified during the merging of genotype files using PLINK v1.9; between-arrays minor allele frequencies differences for overlapping markers were also examined. Final genotype concordance for the four individuals genotyped on all arrays was high (0.999987 for the CNV370v3-Quad-HumanOmniExpressExome and 0.999992 for the HumanExome-HumanOmniExpressExome). Ancestry outliers were detected using multidimensional scaling of distances derived from pairwise kinship coefficients within the KING 1.4 software, leaving 981 RRD Scottish cases for analysis. The final genotype dataset comprises 210071 autosomal variants with MAF > 1%, Hardy-Weinberg equilibrium p-value greater than 1 x $10^{-6}$ and a combined call rate over 99% for all but 10 samples for which exome chip data failed QC.

**Moorfields RRD collection**

Quality controls performed in genome studio and on the exported plink files were as described for the Scottish RRD samples. 1184 samples passed quality controls, with a combined call rate over 99% for markers that included 471,029 autosomal markers with MAF > 1%. N=491 of the 1,184 samples had previously been genotyped for an exome variants analysis using the Illumina HumanExome-12v1-1_A array allowing to check genotyping quality: the concordance for the 18,057 variants with MAF > 1% genotyped on two platforms was 0.999943.

**GS:SFHS**

9,761 samples genotyped using the Illumina HumanOmniExpressExome-8 v1.0 BeadChip were used to provided population matched control for the Scottish RRD analysis. Data quality controls for this dataset were those described in [10] for the OMNI part of the chip array and made use of cluster files and best practices from the CHARGE consortium [11] for the exome part of the chip. GS:SFHS participants identified in [10] to have some Asian or African ancestry were not included but the subgroup with Italian ancestry was kept. In total, 823,358 variants passed genotyping quality control (including N=580,468 autosomal variants with a MAF greater than 1%).

**S5 Text. Imputation to the HRC for the clinical case-control datasets**

As the cases and controls were genotyped separately and on different platforms, extra care was applied. On genotyped data, the consistency of strand calling in cases and controls was checked using –flip-scan option in PLINK, and a case-control analysis was performed on the intercept of datasets using the –assoc option to individually check in genome studio the genotype clusters corresponding to top associated variants. A round of checks was performed prior to phasing and imputation using a script written by W. Rayner available at http://www.well.ox.ac.uk/~wrayner/tools/ : variants with an allele frequency difference larger than 10% compared to the imputation reference dataset (HRC.r1-1) were removed and flagged strand, chromosome , position and naming differences with the HRC reference dataset were acted upon.

**Scottish RRD-GS:SFHS controls**

The intercept strategy described in [12], whereby imputation is performed using the intersection of genotyped variants, was applied. The pre-imputation intercept dataset comprises 193,665 autosomal variants with a MAF>1%. Seven cases in the Scottish RRD study are also GS:SFHS participants, based on their 100% genomic sharing, and their genotypic concordance was high, 0.999993. Multidimensional scaling of distances derived from pairwise kinship coefficients in the intercept dataset, using a LD pruned marker set of N= 77,285 variants, was performed using the KING 1.4 software and showed good ancestry clustering overlap between the two sets (8 Fig). LD pruning of variants was performed in PLINK using the --indep-pairwise 50 5 0.2 argument.

The data was phased using SHAPEIT2 v2r837 [13] with the duoHMM option to use pedigree information to refine phasing [14], and imputation using the HRC reference panel (HRC.r1-1) [8] was performed on the Sanger Imputation Server. The imputed dataset was filtered out for monogenic and low imputation quality variants (INFO < 0.4) using BCFTOOLS 1.3 (http://samtools.github.io/bcftools/). The association analysis was performed further restricting to the 1,136,421 variants with high imputation quality (INFO ≥0.8) and MAF > 1%.

**Moorfield RRD-UK Biobank London controls**

The intercept of the variants passing QC on the Illumina GSA array used to genotype the RRD cases and the Affymetrix array variants available in the UK Biobank controls amounted to N=206,666 autosomal variants with MAF > 1% but the distribution of MAF was clearly shifted towards lower frequency variants (median MAF=0.034 versus 0.24

in the Scottish intercept). Merging of the case and control datasets was therefore performed post imputation [15]. Multidimensional scaling of distances derived from pairwise kinship coefficients in the genotype intercept dataset, using a LD pruned marker set of N= 138,791 variants, showed good ancestry clustering overlap between the two sets (S8 Fig).

A total of 469,574 genotyped variants with MAF > 1% were used as scafold for the HRC imputation of the cases which was performed on the Sanger imputation server after phasing of genotypes using SHAPEIT2 with the duoHMM option. The imputed dataset was filtered out for monogenic and low imputation quality (R2< 0.4) using BCFTOOLS 1.3. Files in vcf format were converted to plink format and merging with the UK Biobank London control subset was performed using plink, leaving N=16,056,823 variants.

A first pass of association uncovered some multiallelic variants not well annotated: for example variant chr3: 16136844 (rs11710284) in the UK Biobank imputation has annotated alleles T and C the latter with AF ~50%, while the same variant, same alleles, has a MAF of 5% in the cases imputed from the GSA array, and also around 5% in the HRC reference panel. Look up in the genome Aggregation Database gnomAD (http://gnomad.broadinstitute.org/)[16] shows that this variant is noted as triallelic with alleles T, C and CCT the latter with frequency 4.8% and 41% in European non Finnish populations. Also evident was that other, not all multiallelic, variants included variants with substantial allele frequency differences: e.g some where AF in cases was close to the gnomAD expectation but differed widely from that in UK Biobank (and sometimes HRC).

Further filtering were made using criteria separately for cases and controls: variants with genotyping call rate < 99% were removed (N=1,906,321 variants in the UK

Biobank imputation),as well as those noted multiallelic in gnomAD, with minor allele frequency difference with HRC reference greater or equal to 10% in cases and controls (N= 266 in cases, N=489 in controls), with minor allele frequency difference with that of non- Finnish European populations in gnomAD reference greater or equal to 5% for controls (N=5907 variants). In addition for GWAS analysis, variants with MAF < 1% (N= 8,453,553 in cases, N= 8,317,862 in controls) were excluded, leaving 5,659,590 variants. Following a second pass of association, further variants were removed: those highly associated variants for which the genotype clusters or that of genotyped variants in LD did not appear well called in genome studio (for the cases), and those for which variants in high LD did not show similar high association P-values (cases and controls).

The final association analysis was performed further restricting to variants with high imputation quality ($R2 \geq 0.9$), amounting to N= 4,727,220 variants.

**S1 Fig**. Study design flow chart

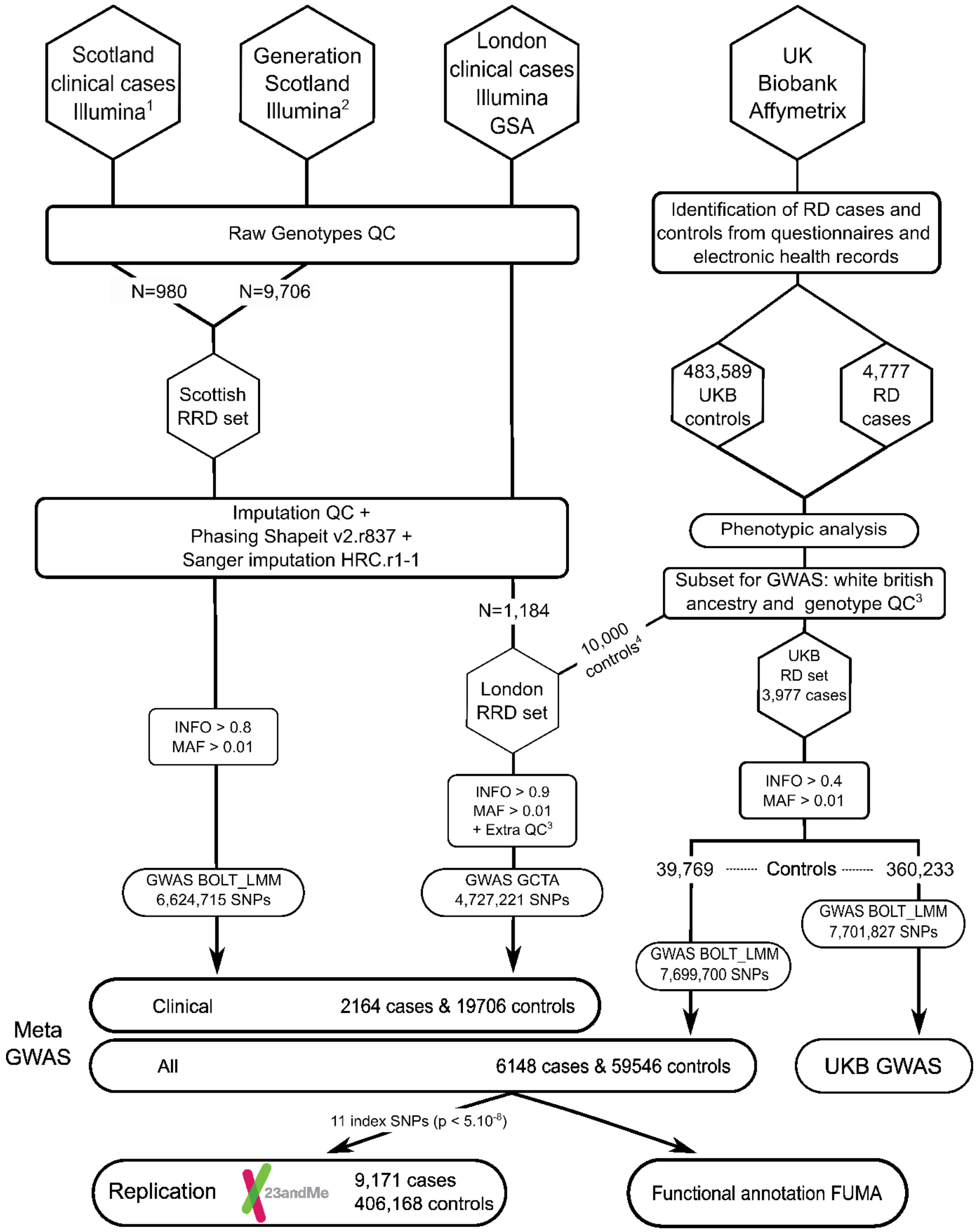


[1] Illumina Quad370 + Exome + OMNI-Express
[2] Illumina OMNI-Express
[3] See Methods section and supplemental information for details
[4] Recruited from London assessment centers

**S2 Fig**. Manhattan and quantile-quantile plots for the UK Biobank retinal detachment primary GWAS (Self report and ICD10 H33 codes or ICD9 316 code; N=3977 cases of white British ancestry; N=360233 controls) using genotypes imputed to the Haplotype Reference Consortium reference.

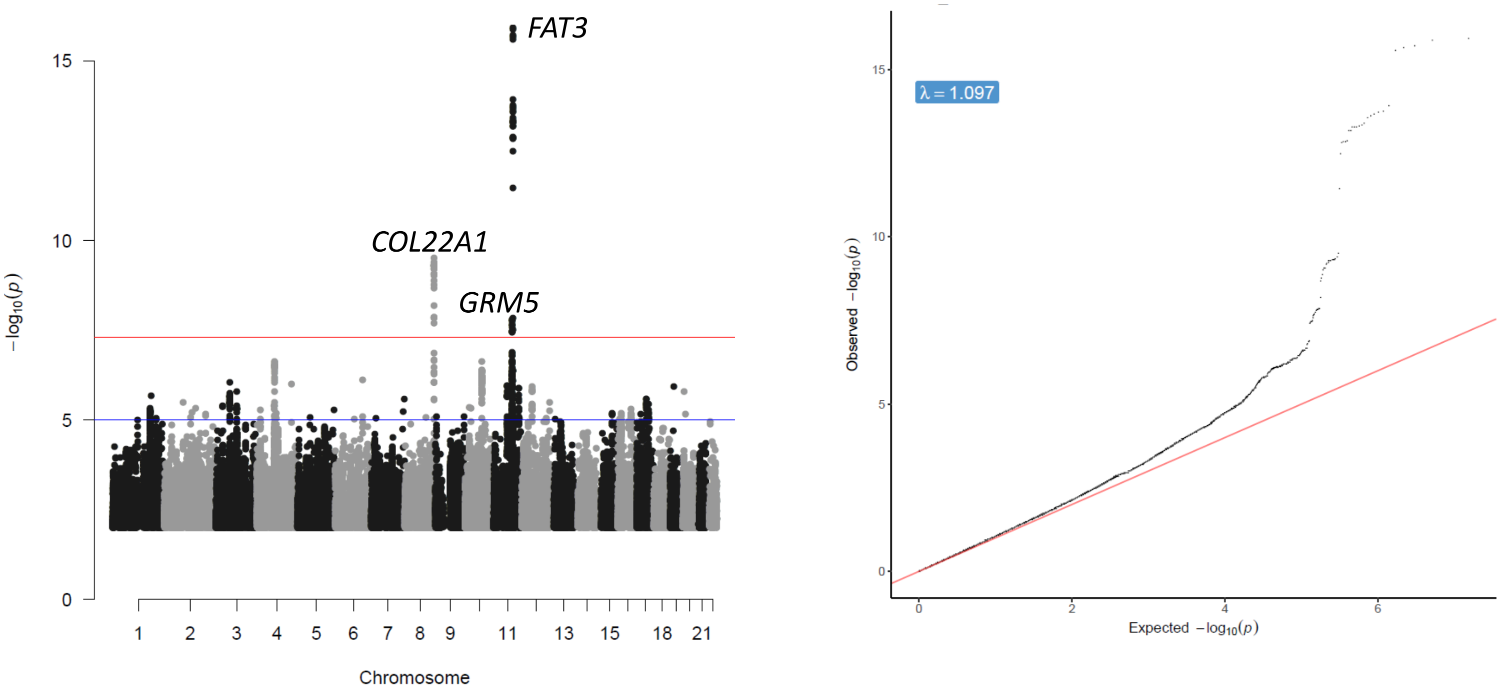

**S3 Fig**. Manhattan plot of the retinal detachment GWAS in UK Biobank (Self report and ICD10 H33 codes or ICD9 316 code; N=3977 cases of white British ancestry) when a 1:3 case to recruitment-centre and sex-matched control ratio is used. Significance thresholds corresponding to P-value=5 x $10^{-8}$ and $10^{-5}$ are represented respectively in red and blue.

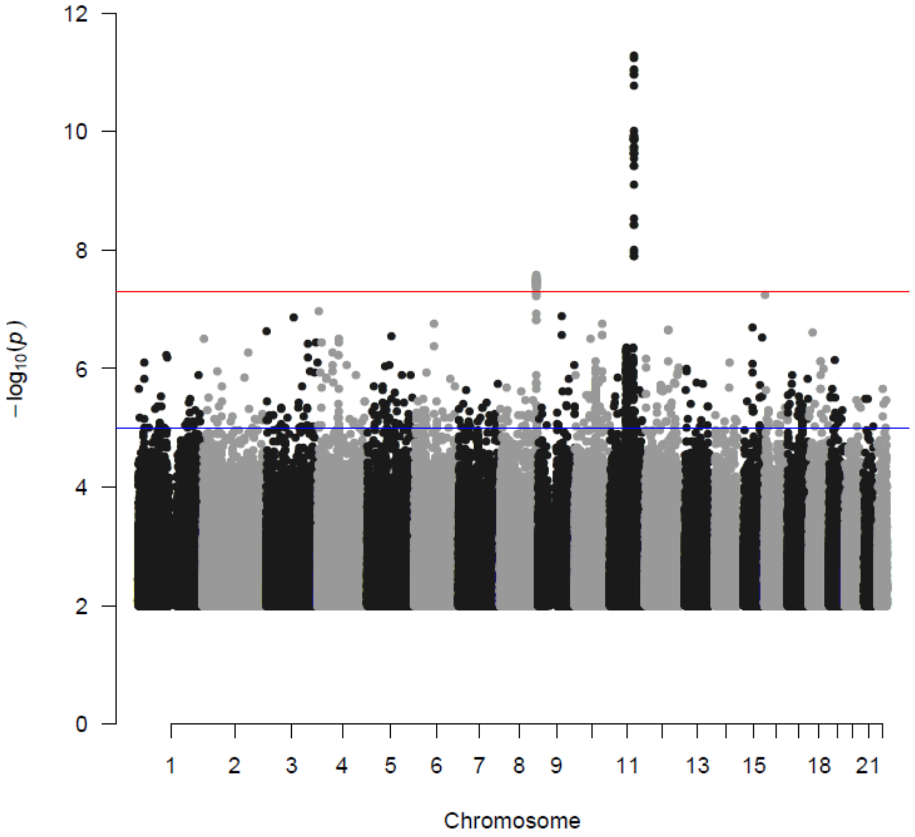

**S4 Fig**. Manhattan plots of GWAS for conditions epidemiologically associated with retinal detachment using the UK Biobank data. Significance thresholds corresponding to P-values of 5 x $10^{-8}$ and $10^{-5}$ are represented respectively by red and blue lines.

A. High Myopia (N=2737 cases N=47635 controls). Genomic inflation factor λ=1.06047, LDscore regression intercept=1.014 (0.006)

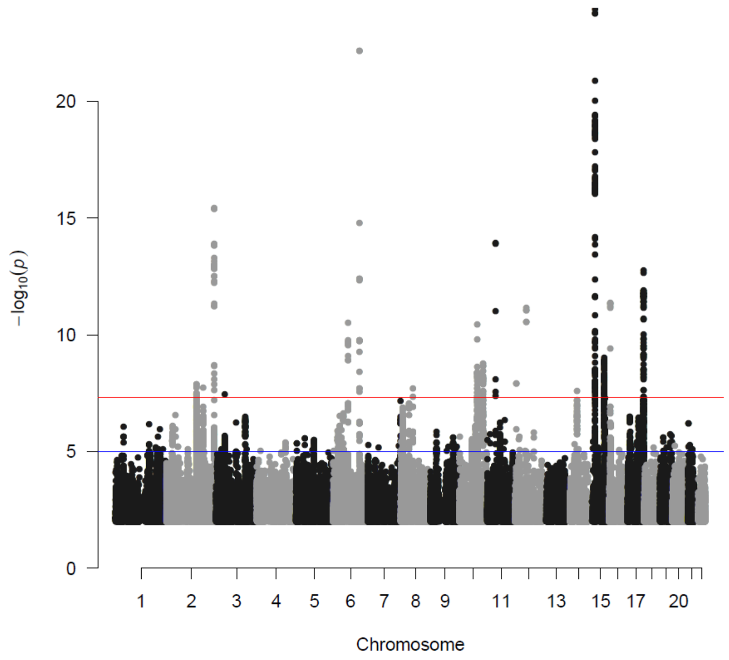


B. History of cataract operation (N=21679 cases N=387283 controls). Genomic inflation factor λ=1.097; LDscore regression intercept=1.019 (0.004)

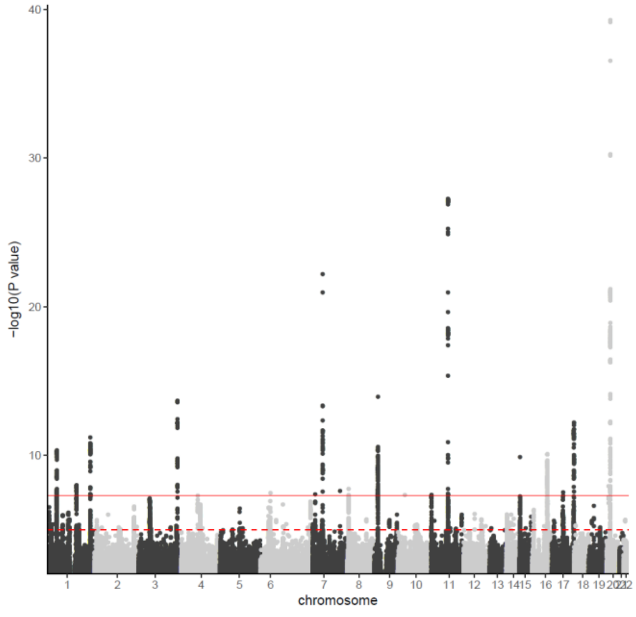

**S5 Fig**. Regional association plots around the RD-SR-ICD UK Biobank loci in epidemiologically related traits

A. Cataract operation regional association plots

COL22A1 GRM5 FAT3

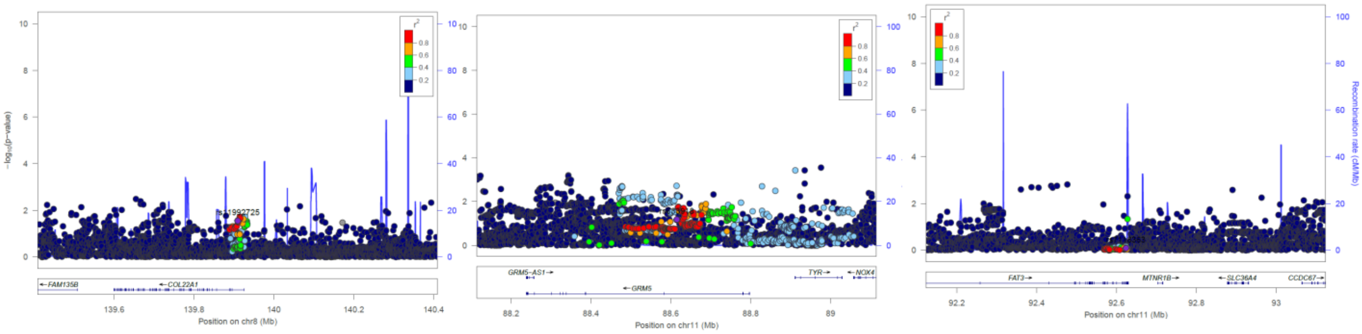

B. High myopia regional association plots

COL22A1 GRM5 FAT3

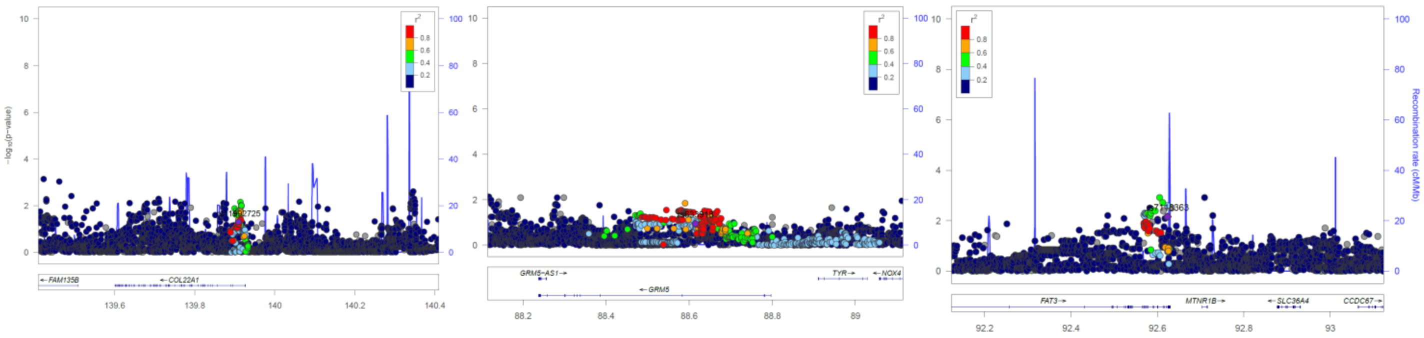

**S6 Fig**. Distributional properties of myopia genetic risk score and retinal detachment status

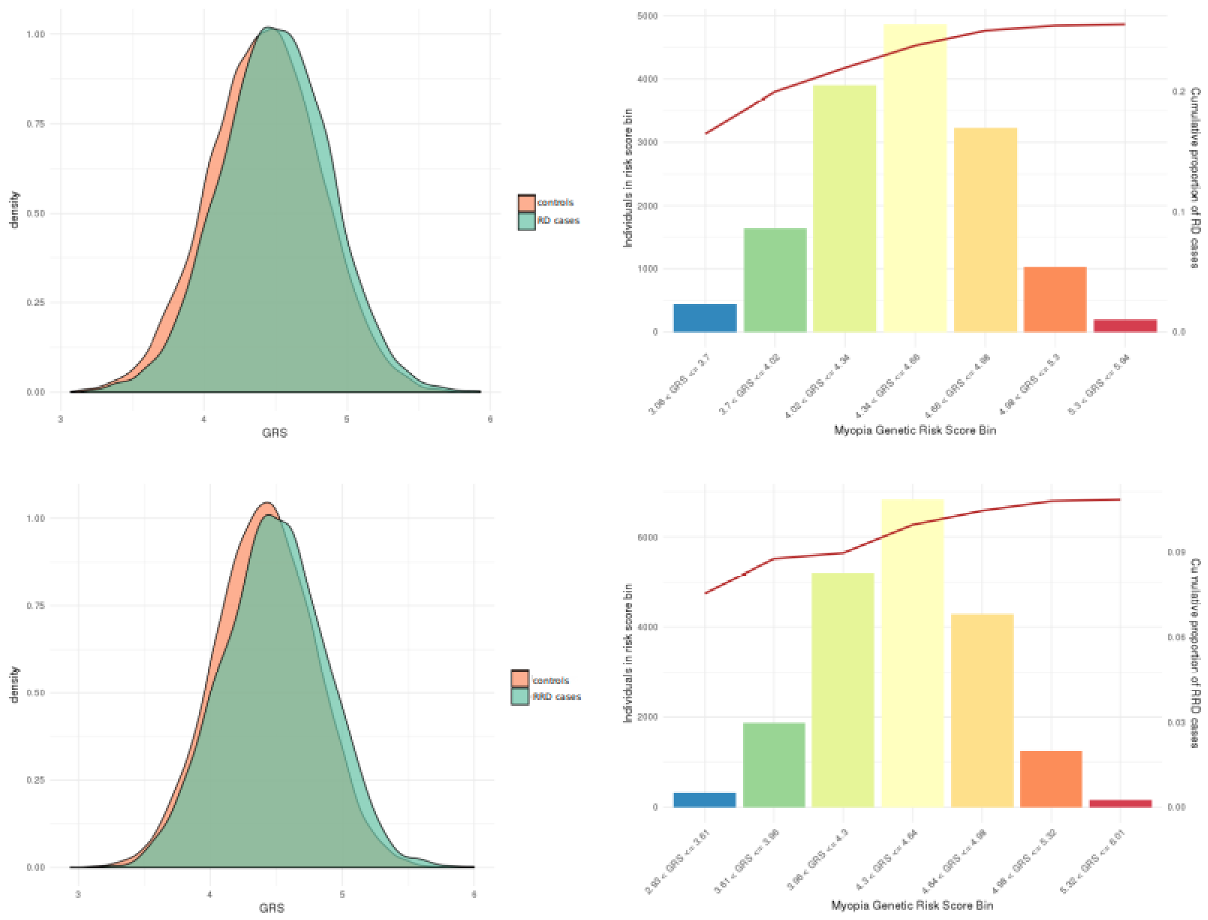


A myopia genetic risk score (GRS) was calculated using summary statistics from 71 independent loci (p-value ≤ $5.10^{-8}$) from a large refractive error meta-analysis[17] with index variants passing quality control in the three investigated cohorts. The distributions of the myopia GRS in the UK Biobank (unrelated only) and the clinically ascertained sets are displayed respectively on the top and bottom plots.

**S7 Fig.** Forest plot for the meta-analysis of effects on retinal detachment of the TYR missense variant rs1042602 (effect reported for the C allele). TE effect size in case control sets with a 1:10 case to control ratio.

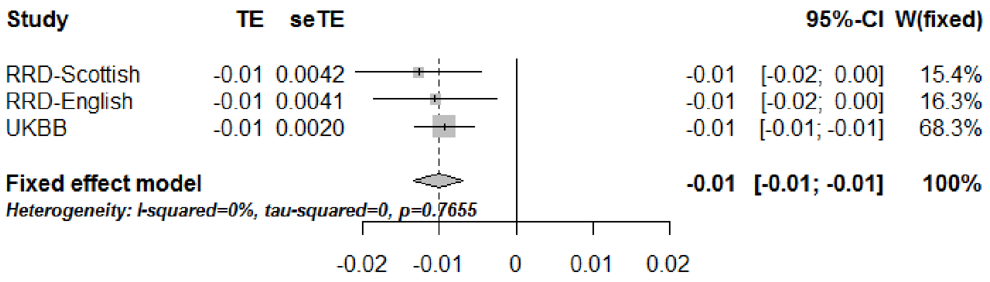

**S8 Fig.** Plots of the first two principal components in multidimensional data reduction of the genotypes available in both clinical cases and controls. The set of variants used for analysis were pruned to remove variants in high linkage disequilibrium. A. Scottish Study RRD cases (N=981) and GS:SFHS controls (N=9754) and B. Moorfield recruited RRD cases and UK Biobank participants recruited in London and defined as of white british ancestry by the UK Biobank central analysis team used as controls. Black circles represent control and red circles cases.

A. Scottish set (N=77285 variants). Individuals with high PC1 scores were shown in previous analysis of GS:SFHS to have Italian heritage.

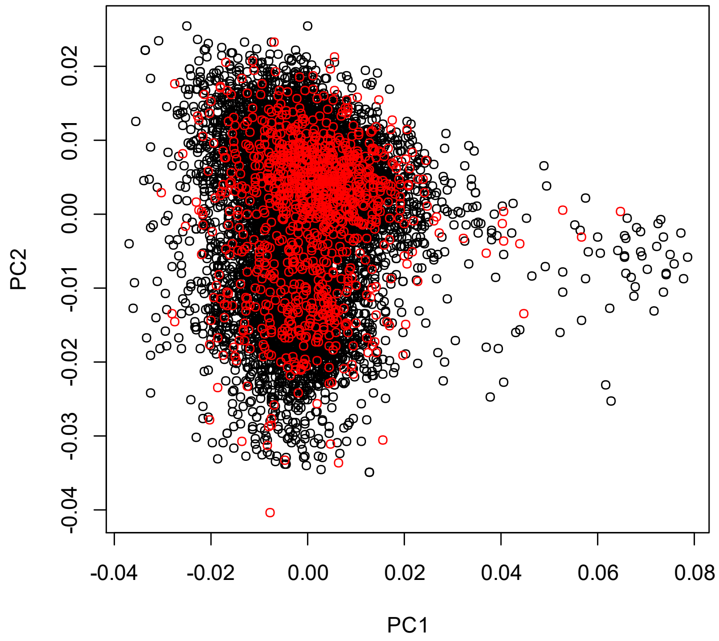


B. London set (N= 138791 variants)

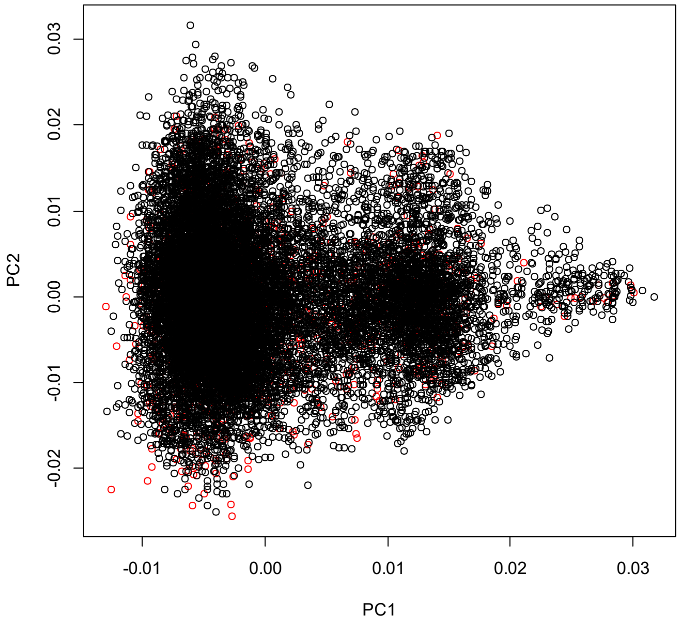

**S1 Table**. Regions associated with retinal detachment in UK Biobank (RD-SR-ICD) using a P-value threshold of $5x10^{-6}$. Genome-wide significant signals are indicated in bold. SNP: index variant in chromosome: position notation with position that of the GRCh37/ hg19 assembly; Name: rs number; N: number of associated variants at P value below threshold within 1Mb; Locus: name of gene if the lead variant within a gene, or if within 20kb of 3'/5' of a gene (closest named) else flanking genes are named separated by --; & indicates locus within which minor allele frequencies of associated variants differ by more than 15%; EA: effect allele; EAF: effect allele frequency (%); Effect: effect size estimate from BOLT-LMM; SE: its standard error; P: P-value.

| SNP | Name | N | Locus | EA | EAF | Effect ($x10^2$) | SE ($x10^2$) | P |
|---|---|---|---|---|---|---|---|---|
| 1:179484502 | rs61827668 | 1 | *AXDND1* | A | 77.2 | 0.132 | 0.029 | 4.8E-06 |
| 1:183786777 | rs114176452 | 1 | *RGL1* | T | 98.9 | -0.585 | 0.124 | 2.1E-06 |
| 2:88415087 | rs1441642 | 1 | 3' *SMYD1* | A | 60.6 | 0.117 | 0.025 | 3.2E-06 |
| 2:147149389 | rs62168444 | 1 | *ZEB2--ACVR2A* | C | 95.7 | -0.276 | 0.061 | 4.7E-06 |
| 3:33174156 | rs1135127 | 3 | *CRTAP* | T | 57.4 | -0.114 | 0.025 | 4.0E-06 |
| 3:66994702 | rs11128322 | 35 | *LRIG1--KBTBD8* | G | 77.1 | -0.144 | 0.029 | 8.9E-07 |
| 3:99914829 | rs80277094 | 7 | 3' *TMEM30C* | G | 81.3 | -0.150 | 0.031 | 1.6E-06 |
| 4:79300406 | rs72657077 | 1 | *FRAS1* | C | 98.0 | -0.402 | 0.087 | 4.1E-06 |
| 4:81267811 | rs556246924 | 1 | *c4orf22* | A | 1.4 | -0.496 | 0.108 | 4.90E-06 |
| 4:81970256 | rs11942256 | 17 | *BMP3*& | G | 96.2 | -0.329 | 0.064 | 2.4E-07 |
| 4:162996109 | rs140229059 | 1 | *FSTL5* | G | 98.9 | -0.608 | 0.125 | 9.9E-07 |
| 6:129820038 | rs12193446 | 1 | *LAMA2* | A | 90.4 | 0.204 | 0.041 | 7.8E-07 |
| 7:157270130 | rs12669108 | 1 | *DNAJB6--PTPRN2* | A | 92.3 | 0.259 | 0.055 | 2.7E-06 |
| **8:139909653** | **rs11992725** | **35** | ***COL22A1*** | **G** | 31.9 | **0.164** | **0.026** | **3.1E-10** |
| 10:20309485 | rs7076757 | 1 | *PLXDC2* | G | 78.7 | 0.140 | 0.031 | 4.4E-06 |
| 10:28064920 | rs113512743 | 1 | *MKX--ARMC4* | C | 96.2 | -0.293 | 0.064 | 4.7E-06 |
| 10:79619461 | rs2579136 | 116 | *DLG5* | G | 70.7 | -0.138 | 0.027 | 2.4E-07 |
| 11:65640562 | rs2303385 | 10 | 5'*EFEMP2* | G | 36.1 | -0.123 | 0.025 | 1.1E-06 |
| **11:88614856** | **rs633918** | **77** | ***GRM5*** | **T** | **61.9** | **0.142** | **0.025** | **1.6E-08** |
| **11:92623608** | **rs7118363** | **32** | ***FAT3*** | **A** | **60.8** | **0.207** | **0.025** | **1.2E-16** |
| 11:120055403 | rs12792820 | 7 | *TRIM29--OAF* | T | 69.4 | -0.128 | 0.026 | 1.3E-06 |
| 12:48374513 | rs1635554 | 10 | *COL2A1* | G | 8.9 | -0.210 | 0.043 | 1.2E-06 |
| 12:132266449 | rs113721561 | 1 | *SFSWAP* | C | 98.3 | -0.446 | 0.096 | 3.2E-06 |
| 17:43991643 | rs9913462 | 9 | *MAPT*& | G | 89.3 | 0.187 | 0.0397 | 2.6E-06 |
| 17:54707917 | rs12939240 | 1 | *ANKNF1--DGKE* | G | 69.2 | 0.122 | 0.026 | 3.7E-06 |
| 19:17089031 | rs73016337 | 1 | *CPAMD8* | A | 98.3 | -0.538 | 0.111 | 1.2E-06 |
| 20:6678883 | rs6085655 | 1 | *FERMT1--BMP2* | G | 98.5 | -0.499 | 0.104 | 1.6E-06 |

**S2 Table**. Regions associated with ICD10 code H33.0 in UK Biobank (N=1380 cases) using a P-value cut-off of $10^{-6}$. SNP: index variant in chromosome: position notation with position that of the GRCh37/ hg19 assembly; Name: rs number; N: number of associated variants at P value below threshold within 1Mb; Locus: name of gene if the lead variant within a gene, or if within 20kb of 3'/5' of a gene (closest named) else flanking genes are named separated by ---; EA: effect allele; EAF: effect allele frequency(%); Effect: effect size estimate from BOLT-LMM; SE: its standard error; P: P-value.

| SNP | Name | N | Locus | EA | EAF | Effect ($x10^2$) | SE ($x10^2$) | P |
|---|---|---|---|---|---|---|---|---|
| 2:114056503 | rs757088184 | 1 | *PAX8--CBWD2* | G | 4.9 | 0.2187 | 0.0427 | 3.0E-07 |
| 3:29055730 | 3:29055730_GTT_G | 1 | *ZCWPW2--RBMS3* | GTT | 92.2 | -0.142 | 0.0285 | 6.8E-07 |
| 8:143817548 | rs149038435 | 9 | *THEM6* | G | 98.9 | -0.36 | 0.0679 | 1.2E-07 |
| 11:92601909 | rs10765567 | 7 | *FAT3* | A | 62.7 | 0.0719 | 0.0146 | 8.6E-07 |

**S3 Table**. Genome-wide significant signals for high myopia in UK Biobank. SNP: index variant in chromosome: position notation with position that of the GRCh37/ hg19 assembly; Name: rs number; N: number of associated variants at P value below threshold within 1Mb; Locus: name of gene if the lead variant within a gene, or if within 20kb of 3'/5' of a gene (closest named), else flanking genes are named separated by ---; EA: effect allele; EAF: effect allele frequency (%); Effect: effect size estimate from BOLT-LMM; SE: its standard error; P: P-value; P_RD : association P-value with RD-SR-ICD in UK Biobank with direction of effect on RD in parentheses for P_RD < 5%.

| SNP | SNP | N | Locus | EA | EAF | Effect | SE | P | P_RD |
|---|---|---|---|---|---|---|---|---|---|
| 2:146929769 | rs570124667 | 119 | ZEB2---ACVR2A | A | 54.7 | 0.00827 | 0.00144 | 1.30E-08 | 0.0042 (+) |
| 2:178661126 | rs7584946 | 32 | PDE11A | T | 57.1 | -0.00792 | 0.00144 | 1.90E-08 | 0.17 |
| 2:233385025 | rs2573210 | 35 | PRSS56 | G | 19.3 | 0.01460 | 0.00181 | 3.90E-16 | 0.89 |
| 3:41199132 | rs773929421 | 1 | CTNNB1 | A | 44.9 | -0.00801 | 0.00147 | 3.50E-08 | 0.38 |
| 6:73643289 | rs7744813 | 27 | KCNQ5 | C | 42.1 | -0.0096 | 0.00145 | 3.20E-11 | 5.5E-05 (-) |
| 6:129820038 | rs12193446 | 21 | LAMA2 | A | 89.9 | 0.02293 | 0.00235 | 7.10E-23 | 7.8E-07 (+) |
| 8:60178580 | rs72621438 | 4 | TOX | C | 64.2 | 0.00814 | 0.00148 | 2.10E-08 | 1.10E-02 (+) |
| 10:86021024 | rs4517452 | 74 | RGR | T | 63.8 | -0.01003 | 0.00151 | 3.70E-11 | 0.016 (-) |
| 10:114773608 | rs55899248 | 36 | TCF7L2 | A | 78 | -0.01034 | 0.00171 | 1.70E-09 | 0.0086 (-) |
| 11:40149305 | rs11602008 | 6 | LRRC4C | A | 83.8 | -0.01476 | 0.00194 | 1.20E-14 | 0.12 |
| 12:56115585 | rs3138142 | 5 | RDH5 | C | 75.2 | 0.01117 | 0.00164 | 6.90E-12 | 4E-05 (+) |
| 12:6954864 | rs5442 | 2 | GNB3 | G | 93.3 | -0.01578 | 0.00284 | 1.20E-08 | 0.45 |
| 14:54672733 | rs547932535 | 25 | BMP4---CDKN3 | C | 69.8 | -0.00872 | 0.00156 | 2.70E-08 | 0.056 |
| 15:35005886 | rs524952 | 141 | GOLGA8B---GJD2 | T | 51.7 | -0.01451 | 0.00142 | 1.20E-24 | 9E-05 (-) |
| 15:79379492 | rs1007365 | 141 | RASGFRF1---ANKRD34C | T | 58 | -0.00871 | 0.00143 | 1.00E-09 | 0.29 |
| 16:7460699 | rs7184522 | 11 | RBFOX1 | A | 64.1 | -0.00997 | 0.00149 | 4.50E-12 | 2.7E-05 (-) |
| 17:79628479 | rs562698611 | 142 | PDE6G | G | 35.8 | 0.01082 | 0.00148 | 1.80E-13 | 1.90E-02 (+) |

**S4 Table**. Genome-wide significant signals for cataract operation in UK Biobank. SNP: lead variant in chromosome: position notation with position that of the GRCh37/ hg19 assembly; Name: rs number; N: number of associated variants at P value below threshold within 1Mb; Locus: name of gene if the lead variant within a gene, or if within 20kb of 3'/5' of a gene (closest named), else flanking genes are named separated by --; EA: effect allele; EAF: effect allele frequency (%); Effect: effect size estimate from BOLT-LMM; SE: its standard error; P: P-value; P_RD : association P-value with RD-SR-ICD in UK Biobank with direction of effect on RD in parentheses for P_RD < 5%.

| SNP | Name | N | Locus | EA | EAF | Effect | SE | P | P_RD |
|---|---|---|---|---|---|---|---|---|---|
| 1:51437247 | rs3176459 | 252 | *CDKN2C* | A | 66.6 | 0.00337 | 0.00052 | 4.70E-11 | 0.12 |
| 1:155076734 | rs7366355 | 50 | 3' *EFNA3* | C | 49.0 | -0.0028 | 0.00049 | 1.00E-08 | 0.58 |
| 1:227172290 | rs12593 | 119 | *ADCK3* | C | 53.3 | -0.00337 | 0.00049 | 5.80E-12 | 0.1 |
| 3:181346937 | rs9842371 | 43 | *SOX2-0T* | C | 64.2 | -0.00388 | 0.00051 | 2.00E-14 | 0.064 |
| 6:32627160 | rs9273404 | 3 | 3' *HLA-DQB1* | A | 53.1 | -0.00293 | 0.00053 | 3.30E-08 | 0.62 |
| 7:11677840 | rs2526099 | 8 | *THSD7A* | A | 52.7 | -0.0027 | 0.0005 | 4.00E-08 | 0.98 |
| 7:46214253 | rs17172647 | 47 | *IGFBP3--LOC730338* | A | 73.5 | -0.00559 | 0.00057 | 6.50E-23 | 0.63 |
| 7:127015083 | rs62621812 | 1 | *ZNF800* | G | 97.9 | -0.00982 | 0.00177 | 2.40E-08 | 0.39 |
| 8:9308156 | rs7828059 | 47 | *LOC157273--TNKS* | T | 48.6 | 0.00277 | 0.00049 | 1.80E-08 | 0.16 |
| 9:22206987 | rs1679013 | 130 | *CDKN2B-AS1--DMRTA1* | C | 54.2 | -0.00397 | 0.00051 | 1.20E-14 | 0.98 |
| 10:29024130 | rs1014607 | 1 | *BAMBI--LINC01517* | G | 69.3 | 0.00291 | 0.00053 | 4.40E-08 | 0.64 |
| 11:213740 | rs12226698 | 27 | *RIC8A* | C | 95.6 | -0.00651 | 0.00119 | 4.70E-08 | 0.64 |
| 11:68941192 | rs79721202 | 158 | 5' *LOC338694* | T | 90.6 | -0.00919 | 0.00084 | 5.40E-28 | 0.33 |
| 15:28230318 | rs1800407 | 66 | *OCA2* | C | 91.4 | -0.00562 | 0.00087 | 1.30E-10 | 0.016 (-) |
| 16:69885166 | rs4985447 | 163 | *WWP2* | G | 49.8 | -0.00317 | 0.00049 | 8.00E-11 | 0.87 |
| 17:40824339 | rs200210041 | 55 | *PLEKHH3* | G | 98.6 | 0.01150 | 0.00208 | 3.00E-08 | 0.85 |
| 17:79564930 | rs71373084 | 86 | *NPLOC4* | C | 64.2 | -0.00358 | 0.00051 | 2.00E-12 | 0.012 (-) |
| 17:81062985 | rs28627461 | 36 | 3' *METRNL* | A | 31.9 | -0.00385 | 0.00054 | 5.80E-13 | 0.39 |
| 20:10683666 | rs2057052 | 4 | *JAG1--LOC101929395* | A | 15.8 | 0.00387 | 0.00067 | 1.20E-08 | 0.73 |
| 20:19457268 | rs4814857 | 146 | *SLC24A3* | A | 84.2 | -0.00893 | 0.00067 | 7.00E-40 | 0.24 |

**S5 Table**. Association look-up for index variants associated with RD-SR-ICD at a P-value cut-off of $5x10^{-6}$. P_HM and P_cataract are the association P-values with respectively high myopia and cataract operation in the UK Biobank dataset (Note that the studies do not have equivalent size and power). Keeping effect allele as defined in S1Table, direction of effect are reported in parentheses when a look-up association is nominally significant.

| SNP | Name | Locus | P_RD | P_HM | P_cataract |
|---|---|---|---|---|---|
| 1:179484502 | rs61827668 | *AXDND1* | 4.8E-06 | 0.79 | 0.92 |
| 1:183786777 | rs114176452 | *RGL1* | 2.1E-06 | 0.12 | 0.055 |
| 2:88415087 | rs1441642 | 3’ *SMYD1* | 3.2E-06 | 0.94 | 0.65 |
| 2:147149389 | rs62168444 | *ZEB2*-- | 4.7E-06 | 8.20E-02 | 0.19 |
| 3:33174156 | rs1135127 | *CRTAP* | 4.0E-06 | 0.19 | 0.23 |
| 3:66994702 | rs11128322 | *LRIG1*-- | 8.9E-07 | 5.50E-02 | 0.75 |
| 3:99914829 | rs80277094 | 3' *TMEM30C* | 1.6E-06 | 0.32 | 0.43 |
| 4:79300406 | rs72657077 | *FRAS1* | 4.1E-06 | 0.29 | 0.12 |
| 4:81765067 | rs10016547 | *c4orf22* (A) | 4.0E-06 (-) | 7.00E-03 (-) | 0.016 (-) |
| 4:81970256 | rs11942256 | *BMP3* (G) | 2.4E-07 (-) | 3.80E-02 (-) | 5.3E-07 (-) |
| 4:162996109 | rs140229059 | *FSTL5* | 9.9E-07 | 0.79 | 0.84 |
| 6:129820038 | rs12193446 | *LAMA2* | 7.8E-07 (+) | 7.10E-23(+) | 0.012 (+) |
| 7:157270130 | rs12669108 | *DNAJB6*--- | 2.7E-06 | 0.41 | 0.54 |
| **8:139909653** | **rs11992725** | ***COL22A1(G)*** | **3.1E-10 (+)** | **4.90E-02(+)** | **0.028 (+)** |
| 10:20309485 | rs7076757 | *PLXDC2* | 4.4E-06 | 0.62 | 0.23 |
| 10:28064920 | rs113512743 | *MKX--ARMC4* | 4.7E-06 | 0.36 | 0.56 |
| 10:79619461 | rs2579136 | *DLG5* (G) | 2.4E-07 (-) | 0.68 | 0.015 (-) |
| 11:65640562 | rs2303385 | *5’EFEMP2* | 1.1E-06 | 6.60E-02 | 0.41 |
| **11:88614856** | **rs633918** | ***GRM5*** | **1.6E-08** | **0.11** | **0.066** |
| **11:92623608** | **rs7118363** | ***FAT3 (A)*** | **1.2E-16 (+)** | **6.90E-03(-)** | **0.81** |
| 11:120055403 | rs12792820 | *TRIM29--OAF* | 1.3E-06 | 0.79 | 0.42 |
| 12:48365265 | rs4760607 | *COL2A1* (G) | 1.2E-06 (-) | 8.70E-02 | 0.011 (-) |
| 12:132266449 | rs113721561 | *SFSWAP* | 3.2E-06 | 0.73 | 0.33 |
| 17:43991643 | rs9913462 | *MAPT* | 2.6E-06 | 0.43 | 0.35 |
| 17:43993376 | rs3785880 (T) | *MAPT* | 2.7E-06 (+) | 0.59 | 0.016 (+) |
| 17:54707917 | rs12939240 | *ANKNF1*-- | 3.7E-06 (+) | 6.90E-06(+) | 4.3E-05(+) |
| 19:17089031 | rs73016337 | *CPAMD8* | 1.2E-06 (-) | 0.17 | 4.0E-04(-) |
| 20:6678883 | rs6085655 | *FERMT1*-- | 1.6E-06 | 0.68 | 0.14 |

**S6 Table**. Regions associated with RRD following meta-analysis of two clinically ascertained cases datasets, using a P-value threshold of $5x10^{-6}$. SNP: lead variant in chromosome: position notation with position that of the GRCh37/ hg19 assembly; N: number of associated variants at P value below threshold within 1Mb; Locus: name of gene if the lead variant within a gene, or if within 20kb of 3'/5' of a gene (closest named), else flanking genes are named separated by --; EA: effect allele; EAF: effect allele frequency (%); $\beta_{RRDmeta}$: meta-analysis inverse variance weighted effect estimate; SE: its standard error; P: P-value; Isq heterogeneity measure (%); P_RD : association P-value with RD-SR-ICD in UK Biobank (case control ratio 1:91) with direction of effect on RD in parentheses.

| SNP | N | Locus | EA | EAF | $\beta_{RRDmeta}$ (direction of effects) | $SE_{RRDmeta}$ | $P_{RRDmeta}$ | Isq | P_RD |
|---|---|---|---|---|---|---|---|---|---|
| 1:219759682 | 9 | 5' *ZC3H11B* | T | 37.2 | 0.014 (++) | 0.0029 | 1.09E-06 | 0 | 7.6e-05 (+) |
| 1:226601738 | 1 | 5' *PARP1* | T | 1.6 | 0.0502 (++) | 0.0109 | 3.90E-06 | 0 | 0.71 (+) |
| 2:158853217 | 1 | *UPP2* | A | 91 | -0.0221 (--) | 0.0048 | 4.92E-06 | 65.8 | 0.04 (-) |
| 2:30414874 | 3 | *YPEL5*--*LBH* | T | 97.8 | -0.0448 (--) | 0.0094 | 1.76E-06 | 35.2 | 0.3 (-) |
| 3:58227720 | 1 | *ABHD6* | T | 96.1 | -0.0325 (--) | 0.0071 | 4.69E-06 | 0 | 0.94 (-) |
| 4:81952637 | 11 | *BMP3* | A | 3 | 0.0376 (++) | 0.0079 | 2.00E-06 | 0 | 7.6e-07 (+) |
| 6:111086833 | 1 | *CDK19* | A | 3.2 | 0.0401 (++) | 0.008 | 5.25E-07 | 56.1 | 0.38 (-) |
| 6:48031994 | 1 | *PTCHD4* | A | 9.2 | 0.0236 (++) | 0.0049 | 1.66E-06 | 0 | 0.31 (+) |
| 6:96002983 | 1 | 5' *MANEA* | A | 1.3 | 0.0573 (++) | 0.0121 | 2.17E-06 | 46.4 | 0.85 (+) |
| 7:24207028 | 12 | *STK31--STY* | A | 88.9 | -0.0221 (--) | 0.0044 | 6.42E-07 | 0 | 0.36 (-) |
| 14:62108781 | 2 | *FLJ22447* | A | 98.5 | -0.0529 (--) | 0.0114 | 3.51E-06 | 0 | 0.97 (+) |
| 18:33466102 | 1 | *GALNT1--C18ORF21* | T | 1.4 | 0.063 (++) | 0.0121 | 1.85E-07 | 0 | 0.32 (+) |

**S7 Table**. Regions associated with retinal detachment in the combined UK Biobank and clinical datasets using a P-value threshold of $10^{-6}$. Genome-wide significant signals are indicated in bold. SNP: lead variant in chromosome: position notation with position that of the GRCh37/ hg19 assembly; N: number of associated variants at P value below threshold within 1Mb; Locus: name of gene if the lead variant within a gene, or if within 20kb of 3'/5' of a gene (closest named), else flanking genes are named separated by --; EA: effect allele; EAF: effect allele frequency (%); Effect: meta-analysis inverse variance weighted effect estimate; SE: its standard error; P: P-value; DIR: the direction of effect for the EA in Scottish RRD, UKBB and English RRD respectively; $P_{het}$ the between studies heterogeneity P-value.

| SNP | N | Locus | EA | EAF % | Effect | SE | P | DIR ($P_{het}$) |
|---|---|---|---|---|---|---|---|---|
| 1:119729001 | 1 | *LOC101929147* | A | 97 | -0.0232 | 0.0047 | 8.80E-07 | --- (0.06) |
| **1:219759682** | **41** | **5' *ZC3H11B*** | **T** | **37.3** | **0.01** | **0.0016** | **1.3E-09** | **+++ (0.23)** |
| 2:186293957 | 1 | *ZNF804A--FSIP2* | T | 92.3 | -0.0154 | 0.003 | 3.46E-07 | --- (0.08) |
| **4:81952637** | **93** | ***BMP3*** | **A** | **2.94** | **0.0317** | **0.0047** | **1.20E-11** | **+++ (0.65)** |
| 4:89753225 | 1 | *FAM13A* | A | 78.7 | 0.0095 | 0.0019 | 8.92E-07 | +++ (0.8) |
| 5:98804516 | 55 | *CHD1--LOC102724855* | A | 1.59 | 0.0337 | 0.0063 | 7.27E-08 | +++ (0.91) |
| 6:129820038 | 1 | *LAMA2* | A | 90.6 | 0.0145 | 0.0027 | 1.26E-07 | +++ (0.32) |
| 7:21768231 | 5 | *DNAH11* | A | 29.7 | -0.0086 | 0.0017 | 7.33E-07 | --- (0.13) |
| 8:22713081 | 1 | *PEBP4* | A | 1.2 | 0.1022 | 0.0201 | 3.80E-07 | +?? (NA) |
| 8:61713231 | 1 | *CHD7* | A | 39.4 | 0.008 | 0.0016 | 8.29E-07 | +++ (0.92) |
| **8:139909653** | **19** | ***COL22A1*** | **A** | **68** | **-0.0101** | **0.0017** | **2.10E-09** | **--- (0.07)** |
| **10:79579222** | **119** | ***DLG5*** | **A** | **29.1** | **0.0102** | **0.0017** | **3.67E-09** | **+++ (0.88)** |
| **10:96038686** | **33** | ***PLCE1*** | **A** | **43.4** | **0.0095** | **0.0016** | **3.64E-09** | **+++ (0.4)** |
| **11:65640906** | **10** | **5' *EFEMP2*** | **T** | **64** | **0.0091** | **0.0016** | **3.50E-08** | **+++ (0.053)** |
| **11:88911696** | **52** | ***TYR*** | **A** | **36.5** | **0.0101** | **0.0016** | **1.01E-09** | **+++ (0.77)** |
| **11:92601909** | **32** | ***FAT3*** | **A** | **62.9** | **0.0143** | **0.0016** | **3.13E-18** | **+++ (0.64)** |
| **11:120030227** | **18** | ***TRIM29--OAF*** | **T** | **68.9** | **-0.0095** | **0.0017** | **2.39E-08** | **--- (0.93)** |
| **12:48404636** | **7** | **5' *COL2A1*** | **T** | **91.3** | **0.0154** | **0.0028** | **4.72E-08** | **+++ (0.2)** |
| 14:43793371 | 3 | *LRFN5--FSCB* | T | 1.1 | 0.089 | 0.0171 | 2.03E-07 | ??+ (NA) |

| | | | | | | | | |
|---|---|---|---|---|---|---|---|---|
| 14:54423053 | 1 | *BMP4* | A | 6.0 | -0.0085 | 0.0016 | 1.31E-07 | --- (0.13) |
| **15:74241624** | **24** | ***LOXL1*** | **A** | **53.1** | **0.0097** | **0.0017** | **3.33E-08** | **++?* (0.29)** |
| 15:93747029 | 19 | *RGMA--MCTP2* | T | 72.1 | 0.009 | 0.0018 | 3.22E-07 | +++ (0.81) |
| 15:96170743 | 14 | *LINC0024--NR2F2-AS1* | T | 45.8 | 0.0079 | 0.0016 | 6.95E-07 | +++ (0.86) |
| 16:62221906 | 1 | *CDH8--CDH11* | A | 43.8 | 0.0088 | 0.0018 | 7.00E-07 | ++? (0.72) |
| 17:18151611 | 1 | *FLII* | A | 33.7 | -0.0092 | 0.0019 | 8.04E-07 | --? (0.63) |
| 20:6678883 | 1 | *CASC20--BMP2* | A | 1.5 | 0.042 | 0.0082 | 3.18E-07 | ?+? (NA) |

* rs12441130 present in the three studies DIR T allele +++ P=3.68 E-08; $P_{het}$ 0.115

**S8 Table**. Notable functional annotations for the lead and tagged variants underlying the 11 identified risk loci for retinal detachment (N=564 variants). Tagged variants in LD with lead variants ($r^2$ >= 0.8) included tested as well as untested variants from the 1000G reference phase 3 within a 250 kb distance of lead variants. Variants with the strongest indications of functionality following annotation by FUMA are shown: with high CADD scores (CADD) which measure how deleterious a variant is predicted to be, with low RegulomeDB scores (RDB – which ranges from 1a for an eQTL variant with evidence of a transcription factor (TF) binding a motif for that TF and a DNase footprint to 7 for variant with no regulatory evidences) or/and with a 15 core chromatin states across 127 tissue/cell types from ENCODE and ROADMAP repositories (minChrState) score lower than 8 which indicates an open chromatin state. Nearest gene (nearestgene) and distance to it (Dist) are based on ANNOVAR annotation using Ensembl gene 85 (GRCh37 human genome assembly). Whether the variant is or not within the physical boundary of the locus is indicated by the indicator posMapFilt (0 for no 1 for yes) and whether it is an eQTL by the indicator poseqtlFilt. eQTL evidences were from GTEx v6, GTEx v7, BIOS QTL and BrainSpan.

**Provided as excel file**

**S9 Table**. Reported trait associations for the lead variants underlying the 11 identified risk loci for retinal detachment. Four GWAS results repositories were used: the GWAS catalogue (r2018-02-06), PhenoScanner (as of March 2018) the Global Biobank Engine (URL: http://gbe.stanford.edu) (GBE) [March 2018] and the GeneATLAS (URL: http://geneatlas.roslin.ed.ac.uk/) [March 2018]. Significance: P-value (P) threshold * $P < 0.05$, ** $P < 1x10^{-5}$, *** $P< 5x10^{-8}$; Categorie: 1 = ocular with no known RD association, 2 = ocular with known retinal detachment risk,3 = retinal detachment, 0 otherwise. For each resource, any phenotype associated at the standard genome-wide significant P-value ($P < 5x10^{-8}$) is listed. If no such trait is reported, the top associated trait is reported, and any ocular trait nominally associated is listed.

**Provided as csv file**

**S10 Table**. Suggestive gene-based associations ($P < 10^{-5}$) performed using MAGMA and the RD meta-analysis summary statistics. Novel RD genes with P-value exceeding the genome-wide significant threshold ($2.74 \times 10^{-6}$ following Bonferroni correction for the N= 18246 genes tested) are highlighted in bold. Positions (start and stop) follow GRCh37 human genome assembly.

| Gene | Chromosome | Start | Stop | P |
|---|---|---|---|---|
| *BMP3* | 4 | 81942119 | 81988685 | 4.40E-10 |
| *LOXL1* | 15 | 74208330 | 74254478 | 1.82E-09 |
| *DLG5* | 10 | 79540549 | 79696378 | 1.85E-09 |
| *GRM5* | 11 | 88227744 | 88809113 | 3.27E-09 |
| *FAT3* | 11 | 92075262 | 92639618 | 6E-09 |
| *TRIM29* | 11 | 119971983 | 120066237 | 5.39E-08 |
| *CTSW* | 11 | 65637280 | 65661212 | 2.77E-07 |
| ***CMSS1*** | 3 | 99526678 | 99907447 | 4.18E-07 |
| *EFEMP2* | 11 | 65623912 | 65651063 | 4.58E-07 |
| *MUS81* | 11 | 65614597 | 65645124 | 5.27E-07 |
| ***BMP2*** | 20 | 6738311 | 6770927 | 1.21E-06 |
| *COL22A1* | 8 | 139590478 | 139936249 | 1.46E-06 |
| ***C16orf45*** | 16 | 15518152 | 15728885 | 2.19E-06 |
| ***DIRC3*** | 2 | 218138742 | 218631316 | 2.32E-06 |
| *HDAC5* | 17 | 42144114 | 42211070 | 3.20E-06 |
| *FILIP1L* | 3 | 99538985 | 99843357 | 3.62E-06 |
| *HAUS8* | 19 | 17150539 | 17196435 | 4.35E-06 |
| *C18orf25* | 18 | 43743519 | 43856949 | 4.72E-06 |
| *TMEM30C* | 3 | 99894531 | 99923035 | 4.96E-06 |
| *LATS1* | 6 | 149969289 | 150049392 | 7.18E-06 |
| *GINM1* | 6 | 149877430 | 149922884 | 7.22E-06 |
| *NKAIN3* | 8 | 63151150 | 63922211 | 7.91E-06 |
| *LYL1* | 19 | 13199847 | 13223975 | 8.18E-06 |
| *ADCY2* | 5 | 7386321 | 7840194 | 9.01E-06 |
| *C4orf22* | 4 | 81246874 | 81894910 | 9.16E-06 |
| *NUP43* | 6 | 150035451 | 150080801 | 9.71E-06 |

**Table S11**. Annotations for the candidate genes selected at each identified RD-associated genetic locus by FUMA (SNP2GENE) or by the gene-based association analysis. SNP2GENE identified genes either by physical mapping to the defined loci or based on eQTL evidences linking an associated variant within the locus to the gene. The maximum CADD score for variants mapping to each gene (posMapMaxCADD), the strength of the eQTL association (eqtlMapminQ) and source of the eQTL studies (eqtlMapts) are indicated. The genes in bold are physically located in the loci. Transcript levels in human ocular tissue are from the Eye Integration v0.62 database where processed deep RNA sequencing data from three ocular tissues are deposited and the Ocular Tissue database (https://genome.uiowa.edu/otdb/) where transcription levels in a wider range of ocular tissues has been assessed using a chip-array. Mouse mutant phenotypes were looked up in the Mouse Genome Informatics resource (http://www.informatics.jax.org/) and the International Mouse Phenotyping Consortium (http://www.mousephenotype.org/) as of March 2018 and human associated conditions in OMIM (https://www.omim.org/)with HPO annotation of commonly associated features from https://rarediseases.info.nih.gov/

**Provided as excel file**

**S12 Table**. Top (adjP < 0.01 ) gene-set or pathway enriched using MAGMA on the RD meta-analysis results. The prioritized genes from the meta-analysis genome-wide significant variants or genes (GWset) and by all variants associated at a lower P-value threshold, $10^{-5}$, (SUGset) were used as input. The GWset includes the 49 genes identified by FUMA SNP2GENE and the 4 additional independent genes detected by gene-based analysis. The SUGset includes 283 genes. GO:Gene ontology; cc: cellular component; mf: molecular functions; bp: biological processes; N:number of genes in gene set.* denotes physically close genes within gene set hence likely biased analysis

| | Category | GeneSet | N | P | adjP | genes |
|---|---|---|---|---|---|---|
| GWset | GO_mf | GO receptor serine threonine kinase binding | 15 | 1.23E-06 | 0.001145 | BMP2:BMP3 |
| | GO_cc | GO basement membrane | 93 | 9.80E-06 | 0.005715 | EFEMP2:COL2A1:LOXL1 |
| | GO_cc | GO extracellular matrix component | 125 | 3.15E-05 | 0.009171 | EFEMP2:COL2A1:LOXL1 |
| SUGset | All Canonical | PID_RXR_VDR_Pathway | 26 | 1.24E-06 | 9.75E-04 | VDR: SREBF1: THRA*: RARA* |
| | GO-bp | Reactome_Nuclear_Receptor_Transcription_Pathway | 48 | 1.47E-06 | 9.75E-04 | VDR: THRA*: NR1D1*: RARA*: NR2F6 |
| | GO-bp | GO_Hormone_Mediated_Signaling_Pathway | 158 | 2.98E-06 | 8.55E-03 | VDR: BMP4: MED24*: THRA*: NR1D1*: RARA*: NR2F6: LATS1 |
| | GO-bp | GO_Steroid_Hormone_Mediated_Signaling_Pathway | 125 | 4.51E-06 | 8.55E-03 | VDR: BMP4: MED24*: THRA*: NR1D1*: RARA*: NR2F6 |
| | GO-bp | GO_Leukocyte_Mediated_Cytotoxicity | 35 | 5.78E-06 | 8.55E-03 | CTSC: RAET1E*: RAET1G*: ULBP2* |
| | GO-cc | GO_Basement_Membrane | 93 | 4.78E-07 | 2.77E-04 | EFEMP2: COL2A1: LOXL1: EFEMP1: FRAS1: LAMA2: ACHE |
| | GO-cc | GO_Extracellular_Matrix_Component | 125 | 4.51E-06 | 1.31E-03 | EFEMP2: COL2A1: LOXL1: EFEMP1: FRAS1: LAMA2: ACHE |
| | GO-cc | GO_Proteinaceous_Extracellular_Matrix | 355 | 1.55E-05 | 2.78E-03 | EFEMP2: COL2A1: BMP4: LOXL1: WNT3: EFEMP1: FRAS1: LAMA2: GPC2: ACHE: COL22A1 |
| | GO-cc | GO_Extracellular_Matrix | 424 | 1.92E-05 | 2.78E-03 | EFEMP2: COL2A1: BMP4: LOXL1: WNT3: EFEMP1: FRAS1: LAMA2: GPC2: PCOLCE: ACHE: COL22A1 |
| | GO-mf | GO_Transcription_Factor_Activity_Direct_Ligant_Regulated_Sequence_Specific_DNA_Binding | 48 | 1.47E-06 | 1.32E-03 | VDR: THRA*: NR1D1*: RARA*: NR2F6* |
| | GO-mf | GO_Receptor_Serine_Threonine_Kinase_Binding | 15 | 3.75E-06 | 1.51E-03 | BMP4: BMP2: BMP3 |
| | GO-mf | GO_Steroid_Hormone_Receptor_Activity | 59 | 5.03E-06 | 1.51E-03 | VDR: THRA*: NR1D1*: RARA*: NR2F6* |

**S13 Table**.Enriched GWAS-based gene sets in the RD meta-analysis results. RD associated variants with a P-value below the threshold of $10^{-5}$ were used as input for FUMA annotation. GWAS for ocular traits are highlighted in bold.

**Provided as excel file**
